# Embryonic stem cell factors DPPA2/4 facilitate a unique chromatin state in non-small cell lung cancer

**DOI:** 10.1101/2025.04.27.650876

**Authors:** Janith A. Seneviratne, Clare L. Crisp, Eleanor Glancy, Natalie Choy, Winnie Tan, Matthew Neve, Melanie Stammers, Tongtong Wang, Ruby Johnstone, Arshnoor Kaur, Katie A. Fennell, Marian Burr, Benjamin L. Parker, Shabih Shakeel, Melanie A. Eckersley-Maslin

**Affiliations:** Peter MacCallum Cancer Centre, Melbourne, Australia; Sir Peter MacCallum Department of Oncology, The University of Melbourne, Australia; Walter and Eliza Hall Institute, 1G Royal Parade, Parkville, VIC 3052 Australia; Department of Medical Biology, The University of Melbourne, Melbourne, VIC 3052 Australia; ARC Centre for Cryo-electron Microscopy of Membrane Proteins, Bio21 Molecular Science and Biotechnology Institute, University of Melbourne, Parkville, Victoria, Australia; Babraham Institute; The John Curtin School of Medical Research, The Australian National University, Canberra, Australian Capital Territory, Australia; Department of Anatomical Pathology, ACT Pathology, Canberra Health Services, Canberra, Australian Capital Territory, Australia; Department of Anatomy and Physiology, The University of Melbourne, Melbourne, Australia; Department of Biochemistry and Pharmacology, The University of Melbourne, Melbourne, VIC 3052, 16 Australia; School of Mathematics and Statistics, The University of Melbourne, Melbourne, VIC 3010, Australia

## Abstract

Embryonic regulators are often re-expressed in cancers, however the functional and molecular significance of this is not always understood. The epigenetic priming factors *Developmental Pluripotency Associated* 2 *and 4* (DPPA2/4) have crucial roles in early development and are implicated in cancer pathogenesis. We reveal in non-small cell lung cancer (NSCLC), DPPA2/4 co-expression is associated with poorly differentiated tumours, impaired patient outcomes and accelerated *in vivo* xenograft tumour growth. Proteomic analyses reveal DPPA2/4 dimerise to enhance their protein stability and binding efficiency to nucleosomes. Our multiomic epigenomic analysis uncovered novel functions for DPPA2/4 in facilitating a unique chromatin landscape in NSCLC cells at active promoters and enhancers. These domains are simultaneously marked by active H3K4me3, PRC1 deposited H2AK119Ub but devoid of PRC2 deposited H3K27me3. This paradoxical uncoupling between the activity of the Polycomb Repressive Complexes is likely driven by H3K27ac at these regions that prevents PRC2 catalytic activity but not recruitment. Our results demonstrate how in NSCLC cells, DPPA2/4 promote a “poised for repression” chromatin state, uncoupling complex recruitment from catalytic activity. Our study highlights how aberrant re-activation of embryonic factors in cancers may take on new functions, promoting tumourigenesis.

## Introduction

Cancers often undergo dedifferentiation and upregulate many transcription and chromatin regulators associated with stem cells and embryonic states. This has been linked with increased plasticity, aggressiveness and ultimately impaired survival outcomes^1–8^. While we often have a strong understanding of the molecular roles and functions for these embryonic factors in developmental and healthy contexts, for many it remains unknown whether they have similar roles in the context of cancer.

Developmental Pluripotency Associated 2 (DPPA2) and 4 (DPPA4) are a pair of functionally intertwined proteins with essential roles during early embryonic development ^9–12^. These non-enzymatic nuclear proteins containing SAP and C-terminal domains predicted to bind nucleic acids and histones respectively^13^. Their heterodimerisation is thought to be important for their function, and in mouse depleting either DPPA2 or DPPA4 in cell lines or embryos has the same phenotypic consequences as depleting both^11,12^. DPPA2/4 are co-expressed predominantly in the germline and early embryo and are silenced by DNA methylation during gastrulation in the mouse^11,14–16^. Despite this tightly restricted expression, maternal and zygotic knockouts of DPPA2/4 survive the period of embryogenesis when they are normally expressed only to succumb shortly after birth^9–12^. This paradoxical uncoupling between the time of expression and phenotype is a defining feature of epigenetic priming factors that facilitate poised chromatin states to enable future gene expression programs ^17^.

In mouse embryonic stem cells (mESCs), DPPA2/4 prime the epigenome to enable future gene expression programs and cellular transitions. Here, DPPA2/4 associate with bivalent promoters that contain both active H3K4me3 and repressive H3K27me3 histone modifications^18,19^. Depleting DPPA2/4 leads to the loss of bivalent chromatin and the accumulation of repressive DNA methylation, preventing these genes from timely activation upon differentiation ^18,19^. Overexpression of DPPA2/4 increases induced pluripotent stem cell reprogramming efficiency ^20^ suggesting these proteins promote cellular plasticity more broadly. However, we have limited understanding of the molecular functions of DPPA2/4 in non-developmental contexts.

In cancer, increased expression of DPPA2 or DPPA4 has been reported in multiple tumour types including lung^21^, ovarian^22,23^, colorectal^24–27^, bladder^27^, prostate^27^ and gastric^28^ cancers, and is associated with reduced overall survival. Human DPPA4 can act as an oncogene and transform fibroblasts ^29^, although the mechanisms by which this occurs remains unclear. Despite their co-expression in embryos^14,15^, DPPA2/4 have not been investigated together within a cancer context. Nor is it known whether the developmental molecular roles for DPPA2/4 are conserved in human cancers, or if the proteins take on new functions in a cancer context.

Here, we investigate the functional and molecular roles for DPPA2/4 in human cancers using multi-omic and biochemical assays to test the hypothesis that DPPA2/4 prime the cancer cell epigenome to facilitate cancer plasticity. Our pan-cancer analysis revealed that co-expression of DPPA2/4 is associated with dedifferentiated phenotypes and impaired overall survival, particularly in non-small cell lung cancers (NSCLC).

Biochemically, the proteins dimerise for their stability and bind nucleosomal DNA at CpG-rich active and bivalent promoters and intergenic enhancers. DPPA2/4 rewire the chromatin landscape in NSCLC cell lines where their binding is associated with active and poised chromatin states. Surprisingly, DPPA2/4 associates with the normally repressive H2AK119Ub modification at active promoters that lack H3K27me3, uncoupling the activity of the two polycomb repressive complexes from one another. Depleting DPPA2/4 in NSCLC cells leads to loss of H2AK119Ub at active H3K4me3-containing promoters and enhancers that were devoid of H3K27me3. Whilst the PRC2 complex was present at many of these unique chromatin regions, high levels of H3K27ac prevent H3K27 methylation, uncoupling complex recruitment from its catalytic activity. Together our study reveals new molecular roles for the embryonic priming factors DPPA2/4 in facilitating a unique H3K4me3-H2AK119Ub-H3K27ac chromatin state in cancer cells.

## Results

### Co-expression of the embryonic regulators DPPA2/4 in cancers correlates with dedifferentiate states and impaired survival

The embryonic epigenetic priming factors DPPA2/4 are uniquely located in tandem on human chromosome 3 and are highly co-expressed in early development^14,15^ (Figure 1A). Upon cell fate establishment DPPA2/4 expression becomes repressed in most post-embryonic tissues (Figure 1B), with the notable exception of the testes (Figure S1A). In contrast, analysis of The Cancer Genome Atlas (TCGA) revealed high expression of DPPA2 and/or DPPA4 in a range of different tumour types (Figure 1C-D, S1B-C). To assess clinical significance of DPPA2/4 co-expression, we performed univariate Cox proportional hazard (CoxPH) modelling. While Testicular Germ-Cell tumours (TGCT) and thymomas (THYM) had the highest proportion of co-expression (Figure 1D), this was not prognostic (Figure 1E, S1D), likely due to DPPA2/4 being expressed in the normal counterparts. In contrast for low grade glioma (LGG), lung adenocarcinoma (LUAD) and lung squamous cell carcinoma (LUSC), co-expression was associated with worse overall patient survival (Hazard ratio (HR):1.63-6.66, p < 0.05) (Figure 1E-F, S1D). We also performed CoxPH modelling when cohorts were subdivided by DPPA2 or DPPA4 expression alone using the same expression cut-offs and observed that only LGG retained prognostic significance, as opposed to LUSC and LUAD cohorts (Figure S1D-F), suggesting that expression of both genes is necessary to confer deleterious phenotypes in non-small cell lung cancer (NSCLC).

**Figure 1:**
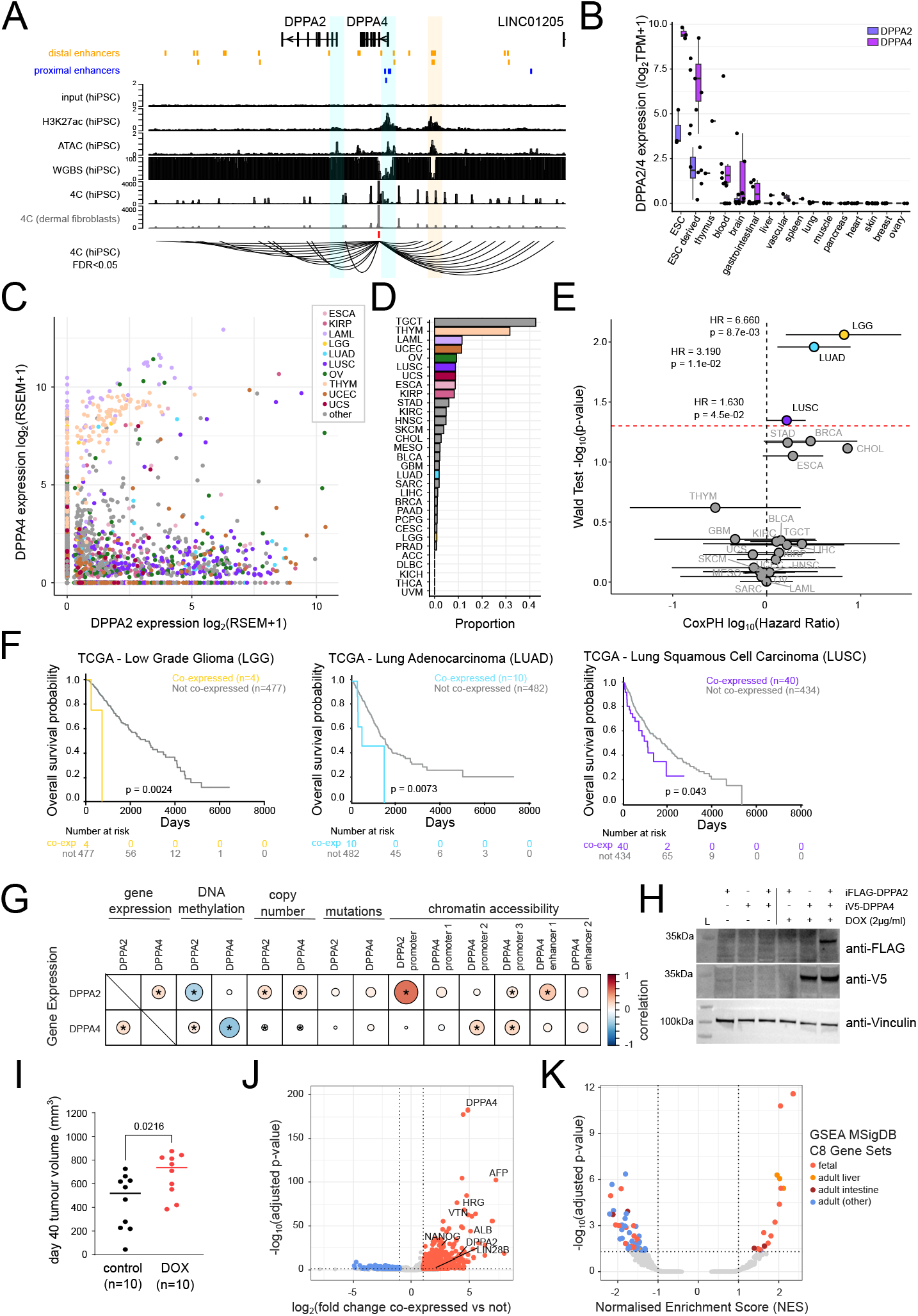
Co-expression of the embryonic regulators DPPA2/4 in cancers correlates with dedifferentiate states and impaired survival. **a)** Overview of the DPPA2/4 genomic locus (hg38; chr3:109238670-109410877) depicting gene structures, ENCODE annotated distal (orange) or proximal (blue) enhancers. Tracks visualise matched genomic input and ChIP-seq for H3K27ac (CPM/bp), chromatin accessibility (ATAC-seq, CPM/bp) and DNA methylation (WGBS, 0-100% DNA methylation) in hIPSCs (reanalysed from GSE145964^71^). Lower tracks visualise 4C-seq following DPPA4 capture in human iPSCs and matched dermal fibroblasts (bait region in red), where axes represent CPM/bp normalised read density (averaged between two replicates) of chimeric reads spanning the capture region and nearby cis-interacting regions (reanalysed from GSE90782^72^). A sashimi track summarises significant cis-interactions (FDR < 0.05) in hiPSC as determined by *r3Cseq*. A putative cluster of enhancers that interact with DPPA4 are highlighted in light orange. **b)** Boxplots of DPPA2/4 expression (transcripts per million, TPM) in a range of human cell line models spanning human development from the epigenetic roadmap consortium. Cell line types ordered by decreasing average DPPA2/4 expression. **c)** Scatter plot visualising DPPA2 (x-axis) and DPPA4 (y-axis) gene expression from RNA-seq across TCGA patients across all cohorts (excluding testicular germ cell tumour (TGCT) cases), selected cohorts are depicted in different colours. Data are represented as log_2_(RSEM+1). **d)** Bar plot displaying proportion of DPPA2/4 co-expressors per cancer cohort (except for the TGCT cohort). **e)** Scatter plot of TCGA cancer cohort Cox proportional hazard modelling (CoxPH) of patient outcomes. Patients in each cohort are subdivided by the co-expression of DPPA2 and DPPA4 as determined by RNA-seq (median of all DPPA2 or DPPA4 expressors (RSEM>0)). The y-axis indicates the statistical significance of CoxPH models (unadjusted p-value of the Wald test for each CoxPH model, −log_10_ transformed), the red line indicates the threshold for p<0.05. The x-axis indicates the hazard ratio (HR) of CoxPH models (log_10_ transformed). Coloured dots indicate those models with HR > 1 and p-value < 0.05, whiskers represent the lower and upper 95% confidence intervals of HRs for each model. **f)** Kaplan-Meier curves demonstrating the stratification of patients into DPPA2/4 co-expressed and low/not-expressed in low grade glioma (LGG, n=481), lung adenocarcinoma (LUAD, n=492) and lung squamous cell carcinoma (LUSC, n=474) TCGA cohorts, respectively. **g)** Dot plot visualising the correlation coefficient of DPPA2/4 gene expression (RNA-seq) correlated with their gene expression (RNA-seq), promoter DNA methylation (methylation arrays), gene copy number (WES), gene somatic mutation (WES) and chromatin accessibility at promoters or distal enhancers (ATAC-seq) in all TCGA cancer cohorts (excluding the TGCT cohort). Biserial correlation is used for mutations, pearson correlation for all others. Significant correlations (BH-adjusted p-value < 0.05) are marked with an asterisk*. **h)** Western blot of whole cell extracts from NCI-H1299 cells following stable integration of dox-inducible Flag-DPPA2 and/or V5-DPPA4 constructs and induction with a vehicle control (H_2_0) or 2μg/mL doxycycline (DOX) for 72h. Blots are anti-Flag, anti-V5 or anti-Vinculin (loading control) with the first lane (L) indicating a molecular ladder with sizes annotated to the left. **i)** Tumour volumes (mm^3^) from Balb/c nude mice xenografted with NCI-H1299 DPPA2/4 double overexpression lines (iFlag-DPPA2 and iV5-DPPA4) at day 40 (last day in which all mice within the study were alive). Lines were pre-treated with a vehicle control (H_2_0) or doxycycline 2μg/mL doxycycline (DOX) for 72h prior to engraftment, and xenografted mice were respectively kept on control or doxycycline feeds to maintain overexpression. **j)** Volcano plot comparing gene expression differences between DPPA2/4 co-expressing vs other NSCLC (LUAD+LUSC) tumours. Differentially expressed genes are those with log_2_FoldChange<-1 (down) or >1 (up) and adjusted p-value<0.05 (EdgeR accounting for tumour type differences (~Type+Group)). **k)** Volcano plot illustrating MSigDB C8 gene sets after a GSEA using all genes ranked by log_2_FC comparing DPPA2/4 co-expressing vs other NSCLC (LUAD+LUSC) tumours. Depleted gene sets are determined as those with NES<-1 and adjusted p-value < 0.05 and enriched gene sets are determined as NES>1 and adjusted p-value < 0.05. Enriched gene sets falling into broad categories are annotated.

We next sought to understand how DPPA2/4 become re-expressed in tumours. In normal somatic tissues, the DPPA2/4 locus is methylated (S1G) suggesting DNA hypomethylation frequently observed in cancer^30^ may lead to DPPA2/4 reactivation. We analysed DPPA2/4 promoter methylation, along with copy number, somatic mutation and chromatin accessibility (Figure S1C), and correlated these with gene expression across all tumour types. TGCT was analysed separately as it arises from a tissue where DPPA2/4 are already highly expressed (Figure 1G, S1A). As expected, expression of DPPA2 and DPPA4 were highly correlated with one another (*r*=0.17, adjP < 0.05) supporting their co-regulation. Consistently, chromatin accessibility at either promoter or at the putative enhancer also weakly correlated with gene expression (Figure 1G).

Overall promoter methylation was highly anti-correlated with gene expression (DPPA2; *r*=-0.301, DPPA4; *r*=-0.39, adjP < 0.05) (Figure 1G, S1H). We also observed modest somatic mutation frequency (9/31 cancer types, 0.6-6.2%) and variable frequency of copy number gains (29/31 cancer types, 0.4 - 66%), however these were only weakly correlated with gene expression (Figure 1G, S1H). These correlations were even more pronounced for TGCT (Figure S1H) reflecting the strong expression of DPPA2/4 in healthy testes (Figure S1A) from which TGCT arises. Taken together these data suggest that DPPA2/4 co-expression in cancer is linked with gene promoter hypomethylation and is associated with worse overall survival outcomes in low-grade glioma and non-small cell lung cancer (NSCLC) patients.

Overexpression of DPPA2 or DPPA4 individually has previously been shown to transform fibroblasts and confer them with oncogenic properties *in vitro* and *in vivo*^29^. To assess whether DPPA2/4 co-expression promote tumour growth in lung cancer we established a doxycycline-inducible system to co-express both DPPA2 and DPPA4 in NCI-H1299 non-small cell lung cancer cells that normally do not express these proteins (Figure 1H) for xenograft experiments. Cells were treated with doxycycline for 3 days prior to subcutaneous engraftment in Balb/c nude mice, and DPPA2/4 co-expression maintained during tumour formation and growth through DOX-chow feed. Both control and DOX-treated tumours were poorly differentiated. At day 40 post-engraftment, when all animals were still alive, there was a significant increase in tumour volume of DOX treated tumours compared to controls (p=0.0216, Figure 1I, S1I-J). This supports a role for DPPA2/4 co-expression in promoting tumour growth in non-small cell lung cancer, ultimately impairing overall survival for these patients.

We next performed differential gene expression analysis to understand the potential consequences of DPPA2/4 co-expression in NSCLC patients (Figure 1E). There were approximately five times more upregulated (n = 1049) that downregulated (n= 206) genes in DPPA2/4 co-expressing NSCLC tumours (|log2FC| > 1, adjP < 0.05). (Figure 1J, Supplemental Table 1). Amongst the upregulated genes were pluripotency markers such as NANOG and LIN28B, as well as several hepatic oncofetal genes such as AFP^5^. Gene set enrichment analyses (GSEA) revealed an enrichment of cellular phenotypes associated with foetal development and regenerative adult tissues, and a depletion of phenotypes associated with non-regenerative adult tissues (Figure 1K, S1I, Supplemental Table 2). This collectively suggests that tumours with DPPA2/4 co-expression recapitulate oncofetal transcriptional programs^5^.

### DPPA2/4 multimerise for stability and nucleosome binding

To study the molecular and cellular functions of DPPA2/4 in a cancer context, we surveyed transcriptomic (Cancer Cell Line Encyclopedia) and proteomic^31^ data, and identified NCI-H661 as one of very few cancer cell lines that co-expressed DPPA2 and DPPA4 at both mRNA and protein levels (Figure S2A-B). NCI-H661 was recently reclassified as a SMARCA4-deficient NSCLC cell line and forms undifferentiated tumours *in vivo*^32^. Using this cell line we generated a series of single and double CRISPR knockout clones (Figure 2A-B, S2C). Notably, the DPPA4 single knockout clones had substantially reduced levels of DPPA2 protein (Figure 2A-B, S1D) despite similar transcript levels (Figure S2C). This suggests that DPPA2 protein requires DPPA4 for its stabilisation. Consistently, overexpression of DPPA2 in NCI-H1299 cells did not result in detectable protein (Figure 1H). In contrast DPPA4 did not appear to require DPPA2 for its stability, and correctly localised to the nucleus in DPPA2 knockout cells (Figure 2A-B). Thus while DPPA2 requires DPPA4 for stability in cells, DPPA4 is stable on its own.

**Figure 2:**
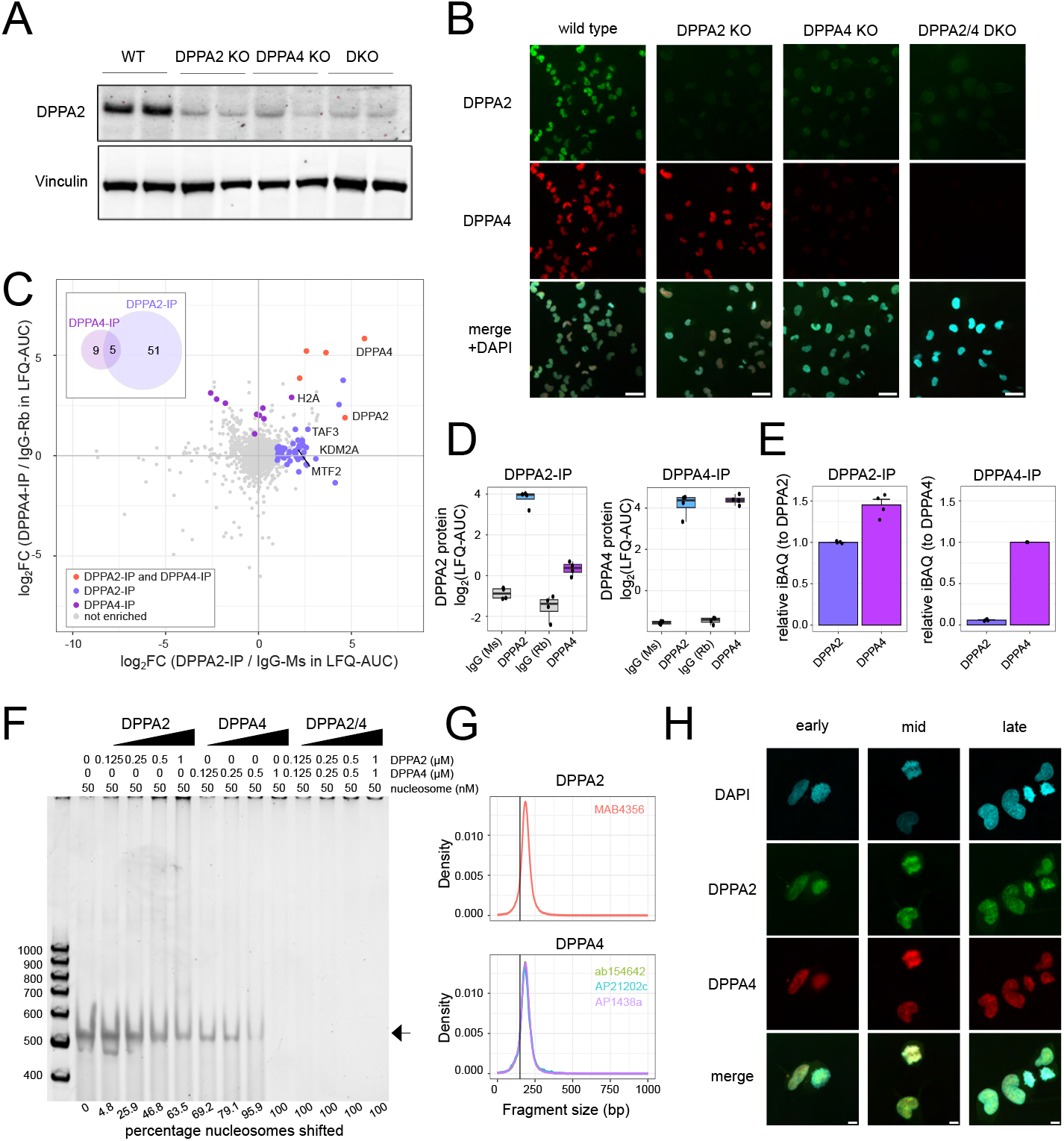
DPPA2/4 multimerise for stability and nucleosome binding. **a)** Western blot of whole cell extracts from NCI-H661 cells following CRISPR-Cas9 mediated knockout of DPPA2 and/or DPPA4 isogenic clones. (WT; wild type, DPPA2 KO; DPPA2 single knockout, DPPA4 KO; DPPA4 single knockout and DKO; DPPA2+4 double knockout). Blots are anti-DPPA2 or anti-Vinculin. **b)** Immunofluorescence images for DPPA2 (green) and DPPA4 (red) in NCI-H661 isogenic knockout clones. Single z-slice representative images are provided for each genotype. Scale bars represent 50μm. **c)** Scatter plot of immunoprecipitation mass spectrometry (IP-MS) results following endogenous DPPA2 or DPPA4 IP from NCI-H661 whole cell lysates (n=4 per condition). Enriched proteins were determined relative to species matched IgG IP control by comparing label-free quantitation (LFQ-AUC) for each protein (log_2_FC ≥ 1 and BH-adjusted p-value < 0.05). The inset Venn diagram highlights the overlap of enriched proteins for DPPA2 and DPPA4. Proteins are coloured based on whether they were enriched in a single IP or both IPs. **d)** Boxplots of enriched DPPA2 and DPPA4 protein levels from IP-MS of DPPA2/4 and matched IgG controls from NCI-H661 whole cell lysates (log_2_LFQ-AUC) (n=4 per condition). **e)** Boxplots of DPPA2 and DPPA4 absolute protein levels relative to the bait (DPPA2/4 stoichiometry) following IP-MS of DPPA2/4 from NCI-H661 whole cell lysates (units are relative iBAQ; intensity-Based Absolute Quantification) (n=4 per condition). **f)** Representative electrophoretic mobility shift assay (EMSA) of unmodified recombinant nucleosomes incubated in the absence/presence of increasing concentrations of recombinant DPPA2-Myc/Flag and/or His-DPPA4 protein. Protein concentrations are denoted atop the gel, and the percentage of nucleosomal DNA bound in each lane is annotated below. Percentage nucleosomes shifted represents the % loss of band compared to input. The experiment was repeated three times. **g)** Density histogram of Enhanced Chromatin Occupancy (EChO) paired-end fragment size analysis from CUT&Run experiments performed using DPPA2 (MAB4356) and DPPA4 (ab154642, AP21202c, AP1438a) antibodies. X-axis represents the mean CUT&Run fragment size at peak foci (peaks called using MACS2 for each CUT&Run relative to a matched IgG control). Vertical line at 150bp represents the cut-off for nucleosomal DNA fragments. **h)** Immunofluorescence imaging of DPPA2 (green) and DPPA4 immunostaining (red) in NCI-H661 parental cells. Representative images are projections (max 5 z-slices) for cells in different stages of mitosis. DAPI stain is shown in blue. Scale bars represent 10μm.

Next, we immunoprecipitated endogenous DPPA2 or DPPA4 in NCI-H661 cells and identified binding partners by mass spectrometry (Figure 2C, S2E-F). As expected, DPPA2 was pulled down by DPPA4 and *vice versa* supporting multimerisation (Figure 2D). To gain insights into potential relative stichometry, intensity based absolute quantification (iBAQ) was performed. Interestingly, when assessing the relative iBAQ values, we observed that in the DPPA2-IP, the ratio between DPPA2 and DPPA4 was >1 (Figure 2E). The converse in the DPPA4-IP was not true, with DPPA2 present at sub-stoichiometric levels relative to DPPA4 (Figure 2C). Thus, while DPPA2 is always found in complex with 1 or more DPPA4 molecules, DPPA4 is stable on its own and may have DPPA2-independent roles. This is despite DPPA2 protein being detected at slightly higher levels compared to DPPA4 (Figure S2C). Consistently, mass photometry analysis showed that only DPPA4 can exist as a monomer *in vitro*, while DPPA2 and DPPA4 form multimers on their own, and higher order complexes when together (Figure S2I-J).

Together our data supports the existence of stable DPPA4 mono/multimers, as well as the canonical DPPA2/4 heterocomplex in NCI-H661 NSCLC cells.

The proteomic analysis revealed 64 proteins that interact with DPPA2 and/or DPPA4 at substoichometric levels (Figure 2C) supporting previous findings in mouse ESCs^19,20^ that these proteins are not stable components of larger complexes. Similar to mouse ESCs ^19,20^ (Figure S2K), we observed association with SETD1A from the MLL/COMPASS-like complex, MTF2 from the Polycomb Repressive Complex 2, as well as the lysine demethylase KDM2A which removes H3K36me2^33^ (Figure 2C, S2L-N). Of interest was a strong association specifically with histone H2A, but not other histones (Figure 2C, S2O-P), suggesting DPPA2/4 may be interacting with nucleosomes rather than naked DNA. To test this, we performed electrophoretic mobility shift assay (EMSA) using recombinant DPPA2 and/or DPPA4 which confirmed DPPA2/4 was able to bind nucleosomes *in vitro* (Figure 2F, S2Q). DPPA4, and to a much lesser extent DPPA2, was also able to bind nucleosomes, albeit not as efficiently as the heterodimer. Supporting nucleosome binding ability, EcHO analysis^34^ of endogenous DPPA2/4 Cut&Run (see below) revealed DPPA2/4 bound fragments were approximately mononucleosomal in length (Figure 2G). Furthermore, and consistent with previous observations in mouse^11,13^, human DPPA2/4 were associated with mitotic chromatin in wild-type NCI-H661 cells (Figure 2B, H). Together these results suggest DPPA4 and DPPA2/4 multimers bind nucleosomes, possibly to facilitate the recruitment and/or stabilisation of chromatin modifying complexes at these loci.

### DPPA2/4 co-locate at active gene regulatory regions marked by H3K4me3, H3K27ac and H2AK119Ub

We next sought to understand where DPPA2/4 bind genome-wide in a cancer context. To this, we performed dual crosslinked chromatin immunoprecipitation of endogenous DPPA2 or DPPA4 followed by high throughput sequencing (ChIP-seq) in NCI-H661 cells, complementing it with native CUT&Run (Figure 3A, S3A-B). This enabled us to determine DPPA2/4 binding sites with high confidence and eliminate any potential transient and/or secondary chromatin binding events that may arise with longer fixation times required for dual crosslinking strategies^35^. There was high concordance between DPPA2 and DPPA4 peaks called using either method supporting their co-occupancy in the genome (Figure S3C). We defined 16220 peaks that were bound by DPPA2 and DPPA4 in all replicates with either ChIP-seq and/or CUT&RUN methodologies (Figure S3B) (termed DPPA2/4 peaks) for subsequent analysis.

**Figure 3:**
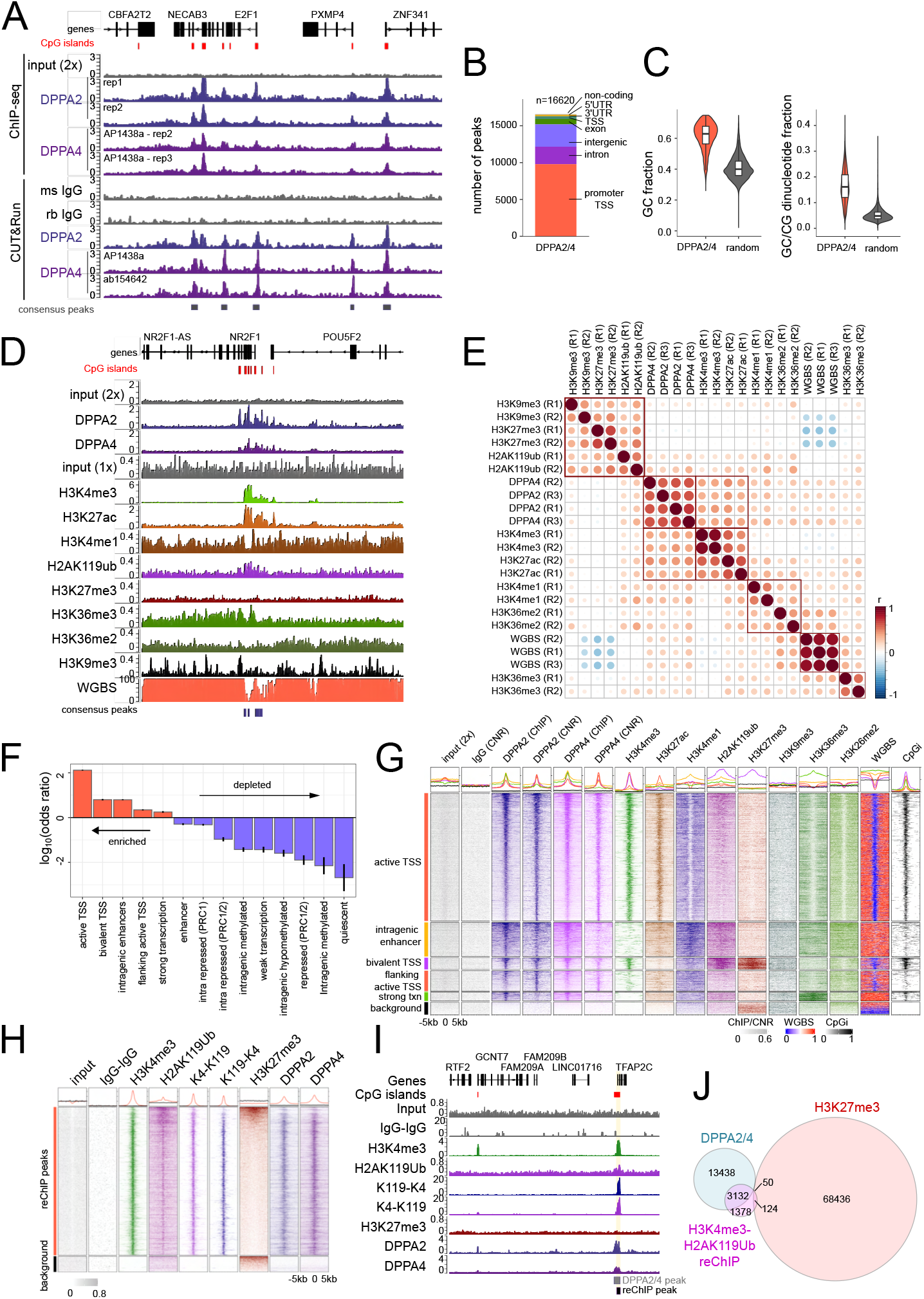
DPPA2/4 co-locate at active gene regulatory regions marked by H3K4me3, H3K27ac and H2AK119Ub. **a)** Genome view of chr 20:33632176-33752019 (hg38) locus. Gene annotation track and CpG islands (red) are shown above. Tracks depict individual replicate tracks of double crosslinked input ChIPs (top) and CUT&Run (bottom). ChIP tracks include control input (n=1, grey), DPPA2 (n=2, blue), DPPA4 (n=2, purple). CUT&Run tracks include mouse and rabbit IgG (n=1 each, grey), DPPA2 (n=1, blue), DPPA4 (n=1, 2 different antibodies, purple). Consensus DPPA2+4 peaks are annotated at the bottom. Scales are in counts per million, per base pair (CPM/bp). **b)** Genomic annotations of DPPA2+4 consensus peaks (n=16620). Promoter-TSS is defined as the TSS+/−1500bp. **c)** G/C content of (left) and GC/CG dinucleotide fraction within (right) of DPPA2+4 consensus peaks (n=16620) compared to size-matched random genomic regions (n=16620). **d)** Genome view of chr 5:93429332-93810092 (hg38) locus. Gene annotation track and CpG islands (red) are shown above. From top to bottom is shown the double crosslinked (2x) input ChIP control (n=1), average DPPA2 (n=2), DPPA4 (n=2) ChIP, single crosslinked input ChIP control (n=1), average H3K4me3 (n=2), H3K27ac (n=2), H3K4me1 (n=2), H2AK119ub (n=2), H3K27me3 (n=2), H3K36me3 (n=2), H3K36me2 (n=2), H3K9me3 (n=2) ChIP and average DNA methylation (WGBS) (n=3). Consensus DPPA2+4 peaks are annotated below. Scales are in CPM/bp for ChIPs and percentage methylated DNA for WGBS (0-100%). **e)** Genome-wide pairwise correlations of ChIP-seq (DPPA2, DPPA4, H3K4me3, H3K27ac, H3K4me1, H3K27me3, H2AK119ub, H3K9me3, H3K36me2 and H3K36me3 fold enrichment over the input control) and WGBS (fraction of methylated DNA) partitioned into 200bp bins genome-wide. Dots are coloured and sized by Pearson correlation coefficients (r) and are ordered by hierarchical clustering with red squares denoting major clusters. **f)** Odds ratio tests of DPPA2+4 consensus regions in association with ChromHMM chromatin states (17 state model, 3 null states (i.e. no overlap with DPPA2+4 consensus regions) excluded. Tests reflect the representation of DPPA2+4 regions in a given chromatin state compared to all other DPPA2+4 regions falling within all other chromatin state annotations. States are ordered by decreasing odds ratio, reflecting enriched chromatin states (log_10_odds ratio>0, red) compared to depleted states (log_10_odds ratio<0, blue). Error bars represent the 95% confidence intervals for hazard ratios, all shown states have an BH-adjusted p-value<0.05. **g)** Heatmaps of genomic regions centred on DPPA2+4 consensus peaks (resized to +/−5kb from the centre of each peak, split into 100 equal windows) overlapping enriched chromatin states: active TSS (n=9676, orange), intragenic enhancer (n=2577, yellow), bivalent TSS (n=890, purple), flanking active TSS (n=1576, orange), strong transcription (n = 633, green) and background regions not bound by DPPA2/4 for comparison (selection, n= 1000, black). Columns represent the double crosslinked input ChIP control (n=1), DPPA2 (n=2) ChIP, DPPA2 (n=1) CUT&Run, DPPA4 (n=2) ChIP, DPPA4 (n=1) CUT&Run, H3K4me3 (n=2), H3K27ac (n=2), H3K4me1 (n=2), H2AK119ub (n=2), H3K27me3 (n=2), H3K9me3 (n=2), H3K36me3 (n=2), H3K36me2 (n=2) ChIPs and DNA methylation (WGBS) (n=3). Last column depicts CpG island density. ChIP and CUT&Run CPM/bp values were scaled per bin across all regions for visualisation. WGBS represents the fraction of methylated DNA. **h)** Heatmaps of genomic regions centred on H3K4me3-H2AK119ub reChIP consensus peaks (resized to +/−5kb from the centre of each peak, split into 100 equal windows) (n = 4663) compared to set of background regions (n =500). Columns represent the double crosslinked input ChIP control (n=1), IgG-IgG reChIP control (n=2), H3K4me3 (n=2), H2AK119ub (n=2), H3K4me3-H2AK119ub reChIP (K4-K119, n=2), H2AK119ub-H3K4me3 reChIP (K119-K4, n=2), H3K27me3 (n=2), DPPA2 (n=2) and DPPA4 (n=2) ChIPs. **i)** Genome view of chr20:56435664-56677328 (hg38) locus. Gene annotation track and CpG islands (red) are shown above. Tracks show input ChIP control (n=1), average reChIP IgG-IgG control (n=2), H3K4me3 (n=2) and H2AK119ub (n=2) ChIP, H3K4me3-H2AK119ub (K4-K119, n=2) and H2AK119ub-H3K4me3 (K119-K4, n=2) reChIP, H3K27me3 (n=2), DPPA2 (n=2) and DPPA4 (n=2) ChIP. Consensus DPPA2/4 (grey) and reChIP peaks (black) are depicted below. Scales are in CPM/bp **j)** Venn diagram showing overlap between DPPA2/4 consensus peaks, H3K4me3-H2AK119ub reChIP peaks (consensus of both K4-K119 and K119-K4 peaks) and H3K27me3 ChIP peaks.

DPPA2/4 bind CG-rich sequences (Figure 3C, S3F) with peaks occurring predominantly at gene promoters (Figure 3B, S3D) of developmental, Wnt signalling and catabolic genes (Figure S3E). To assess the chromatin context for DPPA2/4 binding, we profiled active (H3K4me3, H3K27ac, H3K4me1), repressive (H3K27me3, H2AK119ub, H3K9me3) and transcribing (H3K36me2, H3K36me3) chromatin states, plus whole genome bisulfite sequencing (WGBS) for genome-wide DNA methylation (Figure 3D, S3G-H) in the otherwise uncharacterised HCI-H661 cells. We first partitioned the genome into 200bp bins and hierarchically clustered the signal for each mark against each other (Figure 3E). DPPA2/4 were most highly correlated with active H3K4me3 and H3K27ac marks, and curiously with repressive PRC1-mediated H2AK119ub, but not PRC2-mediated H3K27me3. To explore the chromatin states occupied by DPPA2/4 further, we computed a 17-state ChromHMM hidden Markov model^36,37^ that was able to capture informative combinations of epigenetic modifications (Figure S3I). Odds ratio testing of DPPA2/4 peak overlapping ChromHMM states identified a preference for binding at active (H3K4me3|H3K27ac|H2AK119ub) and bivalent (H3K4me3|H2AK119ub|H3K27me3) TSS states, as well as intragenic enhancers (H3K4me1|H3K27ac) and transcriptional body (H3K36me3) chromatin states (Figure 3F-G). Thus, DPPA2/4 co-occupancy across the genome occurs predominantly at active and poised regulatory regions.

Whilst performing these analyses, we were particularly intrigued by the unusual distribution of H2AK119Ub in NCI-H661 cells. Normally H2AK119Ub, which is deposited by the PRC1 complex, is found together with PRC2-deposited H3K27me3 at repressed gene loci^38,39^. However, we observed many instances, including at DPPA2/4-bound active TSS sites, of H2AK119Ub occurring at H3K4me3 rich regions devoid of H3K27me3 (Figure 3F-G). To eliminate any confounding effects of cellular or allelic heterogeneity in our bulk analyses and confirm true co-occurrence at these sites, we adapted our reChIP protocol^40^ to sequentially immunopreciptate chromatin fragments containing both H3K4me3 and H2AK119Ub. This confirmed the two modifications did indeed co-occur on the same chromatin fragment (Figure 3H-I, S3J), including at active TSS devoid of H3K27me3 (Figure 3J). Notably DPPA2/4 were bound at many of these H3K4me3-H2AK119Ub regions (Figure 3K), suggesting they may have a role in facilitating this unique chromatin structure.

### Loss of DPPA2/4 leads to H2AK119Ub depletion at H2AK119Ub-H3K4me3 promoters

DPPA2/4 are uniquely co-expressed in NCI-H661 compared to other NSCLC cell lines (Figure S2B) which provides an opportunity to investigate how DPPA2/4 may rewire chromatin states in these cells. We reprocessed a comprehensive dataset of 6 chromatin marks that were profiled in 26 NSCLC cell lines^41^ (Figure 4A), which we had also independently assessed in NCI-H661 cells, and generated an integrated 14 state ChromHMM model. When focusing on DPPA2/4 peaks, we noted that NCI-H661 cells clustered away from all other cell lines suggesting the presence of DPPA2/4 induces a unique chromatin state (Figure 4B). This was not observed when a size-matched control set of random genomic regions was compared, arguing against technical or batch differences confounding our analysis (Figure S4A). In NCI-H661 cells, DPPA2/4 bound regions were enriched for active and bivalent TSS and enhancers (Figure 4C) consistent with our observations above (Figure 3F). Strikingly, in NSCLC cells that do not express DPPA2/4, these states were lost with active TSS and enhancers transitioning to a quiescent state, devoid of all 6 chromatin modifications. In contrast, the bivalent TSS transitioned to an active TSS state (Figure 4C). This suggests that DPPA2/4 facilitate these active and poised chromatin states in NCI-H661 cells.

**Figure 4:**
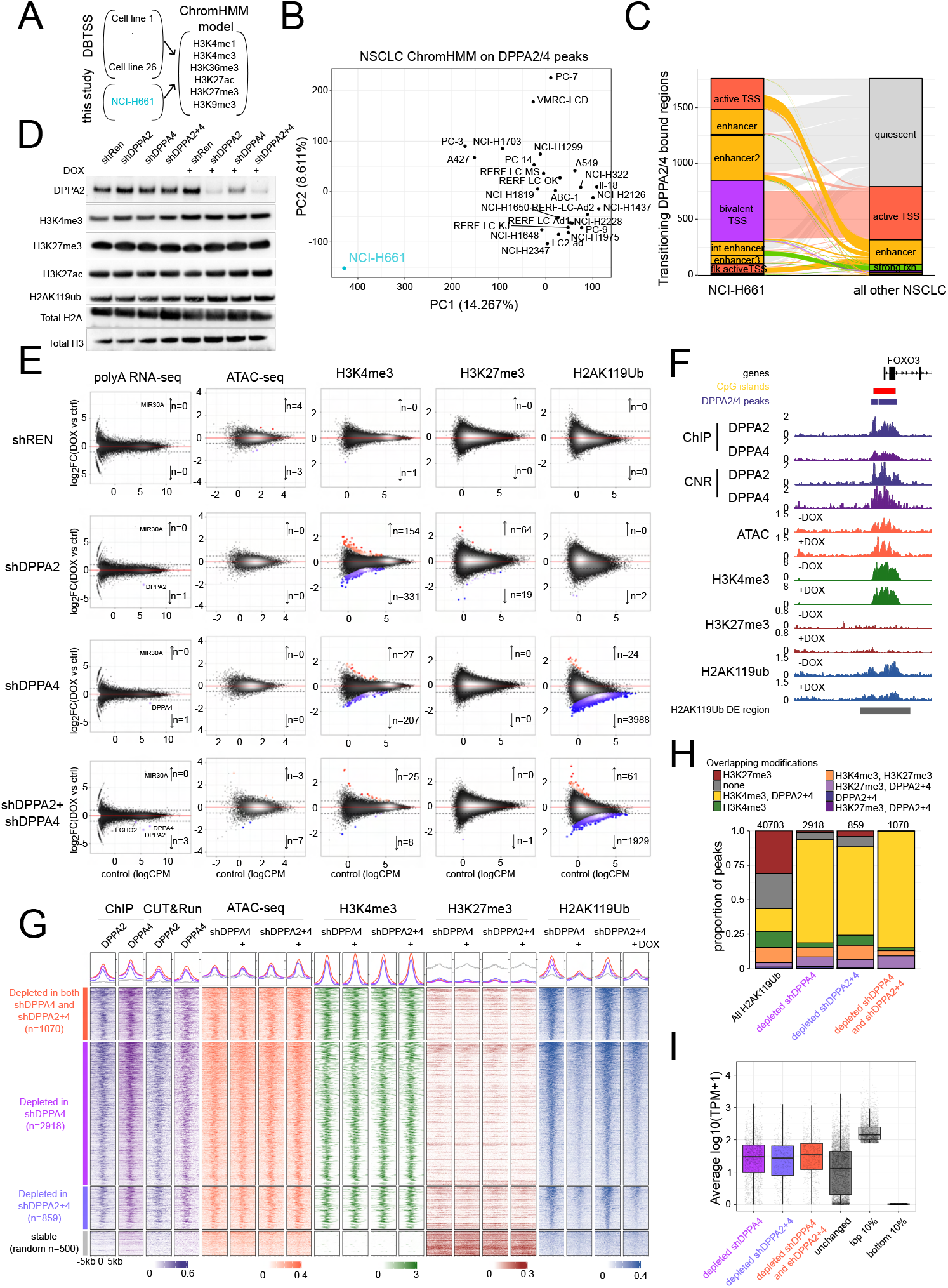
Loss of DPPA2/4 leads to H2AK119Ub depletion at H2AK119Ub-H3K4me3 promoters. **a)** Schematic of data integration between NCI-H661 (this study) and DBTSS (n=26 NSCLC cell lines^41^) to generate a joint 14-state ChromHMM model of all cell lines. **b)** Principle component analysis (PCA) of factorised chromatin states (ChromHMM) at DPPA2/4 peaks (n=16620) across all NSCLC cell lines (n=27). **c)** Alluvial plot of transitioning chromatin states (DPPA2/4 peaks that differ in chromatin state between NCI-H661 cells and all other NSCLC cell lines and are present in at least 75% of other-NSCLC lines). The ribbons between the bars indicate how chromatin states transition between NCI-H661 cells and the majority of other NSCLC cell lines. **d)** Representative western blot of chromatin extracts from stable inducible NCI-H661 cell lines following 7 day shRNA mediated knockdown of DPPA2 and/or DPPA4 (induction with a vehicle control (H_2_O) or 2μg/mL doxycycline (DOX)). Uninduced and induced conditions are provided per cell line (shRen; non-targeting control, shDPPA2; DPPA2 single knockdown, shDPPA4; DPPA4 single knockdown and shDPPA2+4; DPPA2+4 double knockdown). Blots are anti-DPPA2, anti-H3K4me3, anti-H3K27me3, anti-H3K27ac, anti-H2AK119ub, anti-H2A or anti-H3. **e)** Scatter plots of differential RNA-seq and consensus (union overlapping peaks in 2 replicates per condition) ATAC-seq, H3K4me3 ChIP-seq, H3K27me3 ChIP-seq and H2AK119ub ChIP-seq peaks (columns) for the four stable inducible cell lines (first row shREN control, second row shDPPA2, third row shDPPA4 and last row shDPPA2+shDPPA4) comparing 7 day shRNA induction (doxycycline, DOX) to vehicle control (H_2_O). Differentially expressed genes (RNA-seq, FDR<0.05, absolute log_2_FC≥1, edgeR-TMM) or peaks (ATAC-seq, ChIP-seq FDR<0.05, absolute log_2_FC≥0.5, edgeR-TMM) are shown in red (enriched/upregulated) and blue (depleted/downregulated). ATAC-seq and ChIP-seq data were normalised using edgeR-TMM against large 10kb genomic bins genome-wide. Y-axes indicate log_2_FC for DOX vs H_2_O, and x-axes counts per million (logCPM) in the H_2_0 controls. All differential tests represent testing between 3 (RNA-seq) or 2 (ATAC-seq and ChIP-seq) biological replicates per condition. RNA-seq plots also denote miR30A which is part of the shRNA hairpin. **f)** Genome view of chr6:108541111-108580815 (hg38) locus. Gene annotation track, CpG islands (red) and DPPA2/4 peaks (purple) are shown above. Tracks show averaged DPPA2 (n=2) and DPPA4 (n=2) ChIP, DPPA2 (n=1) and DPPA4 (n=1) CUT&Run. Bottom 8 tracks show averaged ATAC-seq, H3K4me3 ChIP-seq, H3K27me3 ChIP-seq and H2AK119ub ChIP-seq for control vs DOX induced shDPPA2+shRPPA4 cells (n=2 each for H_2_O and DOX). Differential H2AK119ub peaks are depicted below. Scales are all CPM/bp. **g)** Heatmaps of genomic regions centred on depleted H2AK119ub peaks (top 3 row groups, peach=depleted in both shDPPA4 and shDPPA2+4 n=1070; purple depleted in shDPPA4 n=2918; blue depleted in shDPPA2+4 n=859) vs stable random regions (bottom group of rows, black, n=500). Regions have been resized to +/−5kb from the centre of each peak, split into 100 equal windows. Columns represent average DPPA2 (n=2) and DPPA4 (n=2) ChIP, DPPA2 (n=1) and DPPA4 (n=1) CUT&Run in parental cells (scaled CPM/bp values across all regions for visualisation), and averaged ATAC-seq, H3K4me3, H3K27me3 and H2AK119ub ChIP-seq for shDPPA4 and shDPPA2+4 (n=2 each for H_2_O and DOX) (unscaled CPM/bp). **h)** Proportional bar plot of overlapping histone modifications (H3K4me3, H3K27me3) and/or DPPA2+4 binding in all (left column) or subsets (right 3 columns) of H2AK119ub regions depleted in shDPPA4 (second column), shDPPA2+4 (third column) or both shDPPA4 and shDPPA2+4 (right column). Total number of peaks per category are annotated atop each bar. **i)** Boxplots of averaged RNA-seq data from parental NCI-H661 cells (n=3) across different categories of H2AK119ub marked promoters, as well as the top 10% and bottom 10% of all expressed genes as controls. Units are in transcripts per million (log10TPM+1).

To test this experimentally we established both siRNA and inducible shRNA knockdown models for either DPPA2, DPPA4 or both DPPA2/4 in the NCI-H661 cell line (Figure S4B-C). This enables the more immediate effects of DPPA2/4 loss to be assessed while avoiding potential confounding clonal effects observed with CRISPR knockout lines^42,43^. Using this approach, we were able to deplete DPPA2 and/or DPPA4 in the cells at both RNA and protein levels (Figure 4D, S4B-C). Similar to above, DPPA4 depletion by shRNA resulted in reduced DPPA2 protein levels (Figure 4D), consistent with the requirement for DPPA4 for a stable functional complex.

We next profiled how DPPA2/4 loss affects transcription and chromatin states in NCI-H661 cells. There were essentially no transcriptional changes observed following DPPA2/4 depletion by shRNA (Figure 4E) or siRNA (Figure S3D-F). Epigenetic priming factors, such as DPPA2/4, often do not directly regulate gene transcription rather they facilitate poised chromatin states^17^. We therefore assessed epigenomic marks that DPPA2/4 are known to regulate in mESCs^18,19^. Surprisingly, there were also no changes in chromatin accessibility nor the canonical bivalent histone modifications H3K4me3 and H3K27me3 following DPPA2/4 shRNA depletion (Figure 4E). Furthermore, DNA methylation landscapes remained largely unchanged when DPPA2/4 were depleted individually or concurrently by siRNAs (Figure S4G-I). Instead, we observed a substantial loss of the PRC1-deposited H2AK119Ub modification in the DPPA4 single and DPPA2/4 double knockdown cells (Figure 4E-F). This was not associated with any changes in global H2AK119Ub (Figure 4D) arguing for locus specific loss rather than a redistribution of the modification.

Closer examination of the H2AK119Ub depleted regions revealed that these were co-occupied by DPPA2/4 and H3K4me3 but lacked H3K27me3 (Figure 4G, S4J). The H2AK119Ub-depleted regions were largely associated with gene promoters associated with developmental processes, Wnt signalling and morphogenesis (Figure S4K-M) that were robustly expressed (Figure 4I). Together these findings point towards a novel role for human DPPA2/4 in cancer cells in facilitating H2AK119Ub at active H3K4me3 regions devoid of H3K27me3.

### H3K27ac prevents H3K27me3 deposition at alternative H3K4me3-H2AK119Ub regions

We lastly sought to further understand these DPPA2/4-dependent H2AK119Ub domains. The absence of H3K27me3 at these sites was of particular interest given the typical co-occurrence of the two polycomb modifications across the genome^38,39^. Two mechanisms could explain this discordance. Firstly, PRC2 may not be effectively recruited and/or displaced these sites. Alternatively, PRC2 may be catalytically inactive or unable to be methylate its substrate because of an alternate competing modification. To understand which of these mechanisms may be involved, we generated ChIP-seq data for SUZ12 (Figure S5A), a core component of PRC2 that helps stabilise the complex^44^, and integrated this with our previous H3K27ac ChIP-seq that we generated as part of our chromatin state modelling (Figure 3F-G).

When analysing the DPPA2/4-dependent H2AK119Ub depleted domains, we noticed that the majority (64%) were also enriched with DPPA2/4, H3K4me3 and H3K27ac (Figure 5A). We termed these regions class I. A smaller (5.3%) set of regions also contained SUZ12 (class II) (Figure 5A-B). Notably, neither of these sets contained H3K27me3 which was very surprising as it suggests an uncoupling between the presence of the PRC2 complex and its catalytic activity at these sites.

**Figure 5:**
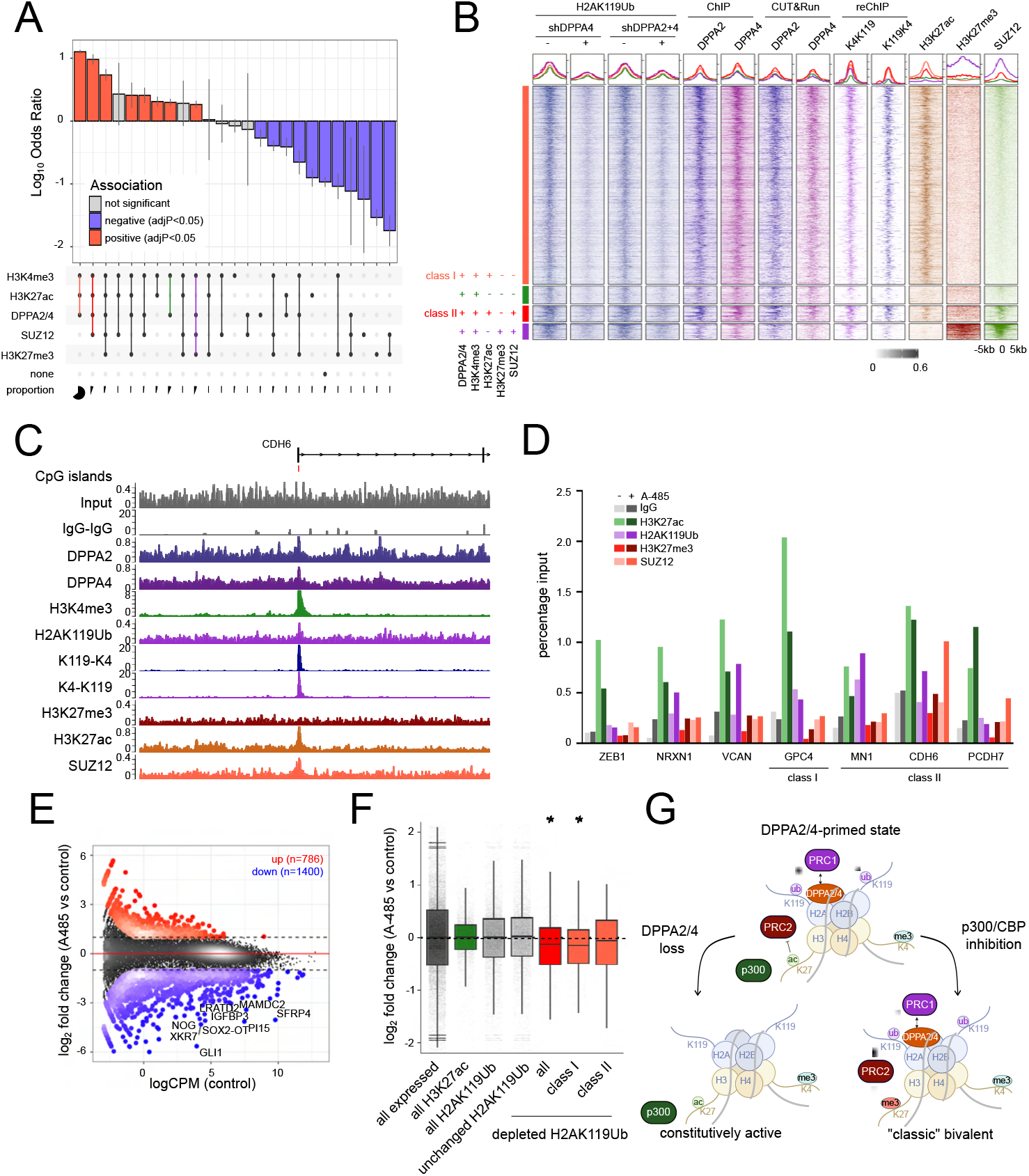
H3K27ac prevents H3K27me3 deposition at alternative H3K4me3-H2AK119Ub regions. **a)** Odds ratio tests of all H2AK119ub depleted regions in association with overlapping H3K4me3, H3K27ac, H3K27me3 and SUZ12 peaks compared to unchanged H2AK119ub regions. Tests reflect the representation of H2AK119ub depleted regions of a particular overlap combination (e.g. H3K4me3+H3K27ac+DPPA2/4) contained within all H2AK119ub depleted regions compared to regions with the same overlap profile falling within unchanged H2AK119ub regions. States are ordered by decreasing odds ratio, reflecting enriched overlaps (log_10_ odds ratio > 0) compared to depleted overlaps (log_10_odds ratio<0). Error bars represent the upper and lower 95% confidence intervals for hazard ratios, coloured overlaps have an BH-adjusted p-value < 0.05. **b)** Heatmaps of genomic regions centred on H2AK119ub depleted peaks grouped by the 4 most abundant overlap categories (resized to +/−5kb from the centre of each peak, split into 100 equal windows). First four columns represent averaged H2AK119ub ChIP in shDPPA4 and shDPPA2+4 control and DOX treated cells (n=2 per condition, unscaled CPM/bp). Remaining columns depict DPPA2 (n=2) and DPPA4 (n=2) ChIP, DPPPA2 and DPPA4 (n=1) CUT&Run, average H3K4me3-H2AK119ub and H2AK119ub-H3K4me3 reChIP (n=2), as well as average H3K27ac (n=2), H3K27me3 (n=2) and SUZ12 (n=1) ChIP in parental cells. Parental cell ChIP and CUT&Run CPM/bp values were scaled per bin across all regions for visualisation. **c)** Genome view of chr5, 31130808, 31270268 (hg38) locus. Gene annotation track and CpG islands (red) are shown above. Tracks (top to bottom) show input ChIP control (n=1), averaged IgG-IgG reChIP control (n=2), average DPPA2 (n=2), DPPA4 (n=2), H3K4me3 (n=2), H2AK119ub (n=2), H2AK119ub-H3K4me3 (n=2), H3K4me3-H2AK119ub (n=2), H3K27me3 (n=2), H3K27ac (n=2) and SUZ12 (n=1) ChIPs/reChIPs. Scales are in CPM/bp. **d)** ChIP-qPCR of H3K27ac (green), H2AK119ub (purple), H3K27me3 (red) and SUZ12 (peach) at selected gene promoters in NCI-H661 cells following treatment with either a vehicle control (DMSO, light) or 10μM A-485 (dark) for 24h (n=1). Values represent the percentage of genomic input used for each ChIP. Class I and II promoters are highlighted. **e)** Scatter plot of RNA-seq data in NCI-H661 cells following treatment with either a vehicle control (DMSO) or 10μM A-485 for 24h (n=3 biological replicates). Differentially expressed genes are highlighted (absolute log_2_FC≥1 and FDR<0.05, edgeR-TMM). Y-axis represents the log_2_FC between A-485 and DMSO control, x-axis indicates the counts per million (logCPM) in the DMSO control. **f)** Boxplots of gene expression changes (log_2_FC) upon A-485 treatment relative to DMSO control across different gene categories: all expressed genes (black, n=29244), all H3K27ac only gene promoters (green, n=3795), all H2AK119ub gene promoters (grey, n=6540), unchanged H2AK119ub gene promoters (grey, n=5877), commonly depleted H2AK119ub gene promoters in shDPPA4 and shDPPA2+4 (red, n=699), class I (n=512) and class II (n=56). Boxes indicate the interquartile ranges, and error bars the upper and lower 95% confidence intervals of all values. **g)** Schematic of the molecular mechanism describing how DPPA2/4 facilitates a *de novo* chromatin state to prime genes for future repression upon H3K27ac removal.

We hypothesised that the presence of H3K27ac at these DPPA2/4-dependent H2AK119Ub domains prevents the recruitment and/or catalytic activity of PRC2 to these sites. To test this, we treated NCI-H661 cells with A-485, a potent inhibitor of p300/CBP that catalyses H3K27ac^45^. As expected, 24 hours of A-485 treatment resulted in reduced H3K27ac levels (Figure 5D). When examining class I or II genes that had the unique H3K4me3-H2AK119Ub-H3K27ac state, the loss of H3K27ac corresponded with an increase in SUZ12 and H3K27me3 at these promoters (Figure 5D). This shift from a more active to repressed chromatin state would likely lead to transcriptional silencing. To this, we performed RNA-sequencing analysis on A-485 treated cells. Consistent with the broad affects of p300/CBP inhibitors, we observed wide-spread transcriptional deregulation with 786 upregulated and 1400 downregulated genes (Figure 5E).

Importantly, those H2AK119Ub enriched promoters that were sensitive to DPPA2/4 loss were repressed upon A-485 treatment. Furthermore, genes associated with both class I and class II domains were also downregulated, although class II did not reach statistical significance, likely due to the smaller number of genes in this category (Figure 5F). This suggests that H3K27ac at these regions maintains the promoter in a transcriptionally active state that is primed for future repression upon removal of the H3K27ac mark.

## Discussion

In this study, we uncover new functions for the embryonic priming factors DPPA2/4 in cancer cells. Our comprehensive pan-cancer analysis reveals how upregulation of DPPA2/4, likely through promoter DNA demethylation, is associated with poorly differentiated phenotypes and worse outcomes, particularly for NSCLC patients.

DPPA2/4 dimerise for their stability and nucleosome binding at CpG rich promoters and enhancers and are required to maintain the canonically repressive H2AK119Ub mark at these regulatory regions. Interestingly, these regions lack detectable H3K27me3, even though PRC2 is able to be recruited in many instances. This uncoupling between complex recruitment and function is likely due to the presence of H3K27ac. Inhibiting the p300/CBP complex reduces acetylation levels allowing PRC2 to rapidly catalyse H3K27me3, resulting in gene repression.

Our study points to novel roles for DPPA2/4 outside of early development, in poorly differentiated cancers. There have been numerous reports of DPPA2 or DPPA4 upregulation in a range of tumour types^21–28^, however the majority of these studies assessed the proteins in isolation and had limited mechanistic follow-up investigations. Our systematic pan-cancer analysis revealed that co-expression of DPPA2/4 is associated with impaired outcomes and dedifferentiated phenotypes in NSCLC. Consistently, co-expression of DPPA2/4 in NSCLC xenograft models results in a modest increase in tumour growth *in vivo*. Similarly, DPPA2 depletion in testicular germ cell tumours impairs tumour growth^46^, and DPPA4 has been shown to transform fibroblasts^29^.

This prompted a biochemical investigation that uncovered a stability axis between DPPA2 and DPPA4. In cells, DPPA2 protein on its own is unstable with very low levels of protein detected in the cells. DPPA2 contains a SPOP-binding consensus degron can be degraded by the SPOP-CUL3-RBX1 E3 ubiquitin-ligase complex^46^, which is expressed in our NSCLC models. It is possible that this degradation is prevented when DPPA4 is present in the cells as the two proteins multimerise. Consistently, despite a direct interaction between SPOP and DPPA2 in mESCs, depleting SPOP has no effect on DPPA2 protein levels^47^. This is potentially due to the substantial levels of DPPA4 in mESCs^14,15,18–20,48^ that would stabilise DPPA2.

In NSCLC cells we reveal a key role for DPPA2/4 in facilitating H2AK119Ub at thousands of active H3K4me3 promoters and enhancers. H2AK119ub is catalysed by Polycomb Repressive Complex 1 (PRC1) and is conventionally a repressive chromatin mark. It is well documented that H2AK119Ub can recruit the PRC2.2 complex to mediate H3K27me3 deposition, which in turn can recruit canonical PRC1^49–58^. Consequently, H2AK119Ub and H3K27me3 typically co-occur to mediate repression, however H2AK119Ub has been shown in some contexst to promote transcription^59^. We reveal this coupling is disrupted in NCI-H661 cancer cells with H2AK119Ub co-occurring with active-associated H3K4me3 and H3K27ac modifications. This highlights the importance of studying H2AK119Ub landscapes which have not been as extensively profiled as H3K27me3, and not to assume mutual exclusivity between H2AK119Ub and H3K27me3.

H3K27 exists in an equilibrium between acetylated and methylated states. Previous studies have revealed a dynamic to-and-fro between these modifications. Disrupting H3K27me3 results in increased acetylation^60–62^, and similarly H3K27ac depletion leads to H3K27me3 accumulation including at sites that are highly transcriptionally active^63^. Our study points towards a role for DPPA2/4 and H2AK119Ub in facilitating the shift from a H3K27 acetylated to methylated state. The transcriptional repression observed with A-485 treatment was predominant at DPPA2/4-dependent H2AK119Ub promoters, suggesting that DPPA2/4 facilitates the poised H2AK119Ub-H3K4me3-H3K27ac state in NSCLC cells. This chromatin state may be priming genes for future repression through a H3K27 acetylation to methylation switch, and have important roles in dynamic systems such as development^62,64^, growth factor stimulation^64^ or cancer cell immune evasion^65^.

Excitingly, our findings point to an uncoupling between the recruitment of PRC2 to chromatin and its catalytic functions. Many of the H2AK119Ub regions that were dependent on DPPA2/4 were bound by SUZ12, a core component of PRC2, yet lacked detectable H3K27me3. This suggests that whilst PRC2 may be recruited to these H2AK119Ub-H3K4me3 domains, its catalytic activity is blocked. We reveal how acetylation of H3K27 likely prevents the activity of PRC2. Similar separation of function for p300/CBP has been recently described whereby p300/CBP is physically present but lacks catalytic activity at repressed promoters^66^. This highlights the importance of assessing both recruitment and function of chromatin regulators as the two properties can be uncoupled and may be contributing to altered pathologies such as cancer.

Bivalent chromatin was first described almost two decades ago in mESCs^67,68^ and has since been reported in multiple contexts and organisms, including human cancers^69,70^. It is traditionally defined as the co-occurrence of H3K27me3 and H3K4me3 on the same nucleosome or chromatin fragment, however many other chromatin marks, including H2AK119Ub, will also co-occur at these domains^70^. Our work highlights the importance of considering other combinations of seemingly opposing histone modifications as poised chromatin states. Cancer can serve as a useful sandbox for discovery for many of these states which may be harder to detect in other cellular contexts as they may be rare or transient in normal cells. Consequently, it will be important to understand how widespread these alternative poised states are in other cellular contexts.

## Resource availability

### Lead contact

Further information and requests for resources and reagents should be directed to and will be fulfilled by the lead contact, Melanie Eckersley-Maslin.

## Materials availability

All plasmids and cell lines generated in this study are available upon request.

## Data and code availability

- Sequencing data will be made available through GEO upon publication of the peer-reviewed manuscript
- Proteomics data will be made available via PRIDE upon publication of the peer-reviewed manuscript
- Imaging, EMSA and western blot source data will be made available on Mendeley Data upon publication of the peer-reviewed manuscript
- No original code was developed as part of this project

## Acknowledgements

We would like to thank all past and present members of the Eckersley-Maslin laboratory for their feedback throughout the project. We thank Ben Blyth and the Models of Cancer Translational Research Centre for their assistance with *in vivo* experiments. We thank Aled Parry for providing inducible shRNA backbones. We acknowledge the Molecular Genomics Core (RRID:SCR_025695), Research Flow Core (RRID:SCR_025550), Centre for Advanced Histology and Microscopy (RRID:SCR_025432), Genotyping Core (RRID:SCR_025622) and Research Laboratory Support Services (RRID:SCR_025699) at Peter MacCallum Cancer Centre, and the Genomics facility at Babraham Institute for their support throughout this project. We also thank Nicholas Williamson, Ching-Seng Ang, Keshava Datta, Swati Varshney and Michael Leeming for instrument support in the Bio21 Mass Spectrometry and Proteomics Facility at The University of Melbourne. Research in the Eckersley-Maslin laboratory is funded by a Snow Medical Fellowship awarded to M.A.E.-M. and the Lorenzo and Pamela Galli Medical Research Trust. Research in the Shakeel laboratory is supported by an NHMRC Investigator grant (GNT 2016827). Our laboratory is located on the lands of the Wurundjeri people of the Kulin Nation and we pay our respects to their elders, past present and emerging, and recognise their continuing connection to country and community.

## Author contributions

Conceptualisation: J.A.S. and M.A.E.-M.; Methodology: J.A.S. and C.C.C.; Formal Analysis: J.A.S., E.G. and W.T.,; Investigation: J.A.S., C.C.C., N.C., E.G., W.T., M.N., M.S., T.W., R.J., A.K., K.F., and B.P.,; Data Curation: J.A.S, T.W. and B.P.; Data Review: M.B.; Writing – original draft: J.A.S and M.A.E.-M.; Writing – review and Editing: J.A.S. and M.A.E.-M.; Visualisation: J.A.S. and M.A.E.-M.; Supervision: J.A.S, S.S. and M.A.E.-M.; Project Administration and Funding Acquisition: M.A.E.-M.

## Supplemental information

Document S1. Extended Materials and Methods, legends to Extended Figures 1-5,

**Table 1**: Patient survival statistics for DPPA2, DPPA4 and DPPA2/4 co-expressing tumours, relating to Figure 1, Ext Figure 1E-F

**Table 2**: Correlation values, relating to Figure 1G, Ext Fig 1H

**Table 3**: GSEA gene sets enriched in DPPA2/4 co-expressing tumours, relating to Figure 1J, Ext Figure S1K

**Table 4**: Protein quantification and statistics from proteomic experiments relating to Figure 2C, Ext Figure 2G,H,L,M,N,O,P

**Table 5**: Peak tables for all ChIP-seq, CUT&Run and reChIP-seq experiments, as well as DPPA2/4 and reChIP consensus peak sets related to Figure 3/Ext Figure 3

**Table 6**: Table of correlation statistics relating to Figure 3E

**Table 7**: Table of statistics for chromatin state enrichments relating to Figure 3F **Table 8**: shRNA: Annotated differential RNA-seq, ATAC-seq and ChIP-seq statistics relating to Figure 4E.

**Table 9**: H2Aub differential region overlap enrichment statistics relating to Figure 5A

**Table 10**: Differential RNA-seq statistics, relating to Figure 5E

**Table 11:** TCGA RNA-seq differentially expressed genes in LUAD+LUSC, DPPA24 co-expressors vs other patients.

**Table 12:** GO (biological processes) pathway analyses of DPPA2+4 bound promoters related to Ext Figure 3F.

**Table 13:** H661 siRNA differential RNA-seq and WGBS relating to Ext Figure 4D-I

**Table 14:** Motif analyses of H2Aub differential regions using monaLisa

**Table 15:** GO (biological processes) pathway analyses of H2Aub differential regions

## Methods

See supplemental file for detailed experimental and computational methods.

## References

1. Li, J. & Stanger, B. Z. How Tumor Cell Dedifferentiation Drives Immune Evasion and Resistance to Immunotherapy. Cancer Res. 80, 4037–4041 (2020).

2. Boumahdi, S. & Sauvage F. J. de. The great escape: tumour cell plasticity in resistance to targeted therapy. Nat. Rev. Drug Discov. 19, 39–56 (2020).

3. Hanahan, D. Hallmarks of Cancer: New Dimensions. Cancer Discov 12, 31–46 (2022).

4. Aiello, N. M. & Stanger, B. Z. Echoes of the embryo: using the developmental biology toolkit to study cancer. Dis Model Mech 9, 105–14 (2016).

5. Sharma, A., Blériot, C., Currenti, J. & Ginhoux, F. Oncofetal reprogramming in tumour development and progression. Nat Rev Cancer 1–10 (2022) doi:10.1038/s41568-022-00497-8.

6. Hölzel, M., Bovier, A. & Tüting, T. Plasticity of tumour and immune cells: a source of heterogeneity and a cause for therapy resistance? Nat. Rev. Cancer 13, 365–376 (2013).

7. Yuan, S., Norgard, R. J. & Stanger, B. Z. Cellular Plasticity in Cancer. Cancer Discov. 9, 837–851 (2019).

8. Friedmann-Morvinski, D. & Verma, I. M. Dedifferentiation and reprogramming: origins of cancer stem cells. EMBO Rep. 15, 244–253 (2014).

9. Madan, B. et al. The pluripotency-associated gene Dppa4 is dispensable for embryonic stem cell identity and germ cell development but essential for embryogenesis. Mol Cell Biol 29, 3186–203 (2009).

10. Nakamura, T., Nakagawa, M., Ichisaka, T., Shiota, A. & Yamanaka, S. Essential roles of ECAT15-2/Dppa2 in functional lung development. Mol Cell Biol 31, 4366–78 (2011).

11. Kubinyecz, O., Santos, F., Drage, D., Reik, W. & Eckersley-Maslin, M. A. Maternal Dppa2 and Dppa4 are dispensable for zygotic genome activation but important for offspring survival. Development 148, dev200191 (2021).

12. Chen, Z., Xie, Z. & Zhang, Y. DPPA2 and DPPA4 are dispensable for mouse zygotic genome activation and preimplantation development. Development (2021) doi:10.1242/dev.200178.

13. Masaki, H., Nishida, T., Kitajima, S., Asahina, K. & Teraoka, H. Developmental pluripotency-associated 4 (DPPA4) localized in active chromatin inhibits mouse embryonic stem cell differentiation into a primitive ectoderm lineage. J Biol Chem 282, 33034–42 (2007).

14. Maldonado-Saldivia, J. et al. Dppa2 and Dppa4 are closely linked SAP motif genes restricted to pluripotent cells and the germ line. Stem Cells 25, 19–28 (2007).

15. Eckersley-Maslin, M. et al. Dppa2 and Dppa4 directly regulate the Dux-driven zygotic transcriptional program. Gene Dev 33, 194–208 (2019).

16. Monk, M., Hitchins, M. & Hawes, S. Differential expression of the embryo/cancer gene ECSA(DPPA2), the cancer/testis gene BORIS and the pluripotency structural gene OCT4, in human preimplantation development. Mhr Basic Sci Reproductive Medicine 14, 347–55 (2008).

17. Eckersley-Maslin, M. A. Keeping your options open: insights from Dppa2/4 into how epigenetic priming factors promote cell plasticity. Biochem Soc T 48, 2891–2902 (2020).

18. Gretarsson, K. H. & Hackett, J. A. Dppa2 and Dppa4 counteract de novo methylation to establish a permissive epigenome for development. Nat Struct Mol Biol 27, 706–716 (2020).

19. Eckersley-Maslin, M. A. et al. Epigenetic priming by Dppa2 and 4 in pluripotency facilitates multi-lineage commitment. Nat Struct Mol Biol 27, 696–705 (2020).

20. Hernandez, C. et al. Dppa2/4 Facilitate Epigenetic Remodeling during Reprogramming to Pluripotency. Cell Stem Cell 23, 396–411 e8 (2018).

21. John, T. et al. ECSA/DPPA2 is an embryo-cancer antigen that is coexpressed with cancer-testis antigens in non-small cell lung cancer. Clin Cancer Res 14, 3291–8 (2008).

22. Deb, S. et al. ECSA/DPPA-2 Is a novel embryo-derived molecule which may regulate immune targets in gynaecological malignancies. Pathology 42, S59–S59 (2010).

23. Tchabo, N. E. et al. Expression and serum immunoreactivity of developmentally restricted differentiation antigens in epithelial ovarian cancer. Cancer Immun 9, 6 (2009).

24. Ghodsi, M., Jafarian, A. H., Montazer, M., Sadeghian, M. H. & Forghanifard, M. M. Diagnostic clinical relevance of developmental pluripotency-associated 2 (DPPA2) in colorectal cancer. Int J Surg 13, 193–197 (2015).

25. Zhang, M. et al. Developmental pluripotency-associated 4: a novel predictor for prognosis and a potential therapeutic target for colon cancer. J Exp Clin Canc Res 34, 60 (2015).

26. Raeisossadati, R. et al. Aberrant expression of DPPA2 and HIWI genes in colorectal cancer and their impacts on poor prognosis. Tumor Biol 35, 5299–305 (2014).

27. Amini, S., Fathi, F., Mobalegi, J., Sofimajidpour, H. & Ghadimi, T. The expressions of stem cell markers: Oct4, Nanog, Sox2, nucleostemin, Bmi, Zfx, Tcl1, Tbx3, Dppa4, and Esrrb in bladder, colon, and prostate cancer, and certain cancer cell lines. Anat Cell Biology 47, 1–11 (2014).

28. Shabestarian, H. et al. DPPA2 Protein Expression is Associated with Gastric Cancer Metastasis. Asian Pac J Cancer P 16, 8461–5 (2015).

29. Tung, P. Y., Varlakhanova, N. V. & Knoepfler, P. S. Identification of DPPA4 and DPPA2 as a novel family of pluripotency-related oncogenes. Stem Cells 31, 2330–42 (2013).

30. Ehrlich, M. DNA hypomethylation in cancer cells. Epigenomics 1, 239–259 (2009).

31. Gonçalves, E. et al. Pan-cancer proteomic map of 949 human cell lines. Cancer Cell 40, 835–849.e8 (2022).

32. Ng, J. et al. Molecular and Pathologic Characterization of YAP1-Expressing Small Cell Lung Cancer Cell Lines Leads to Reclassification as SMARCA4-Deficient Malignancies. Clin. Cancer Res. 30, OF1–OF13 (2024).

33. Tsukada, Y. et al. Histone demethylation by a family of JmjC domain-containing proteins. Nature 439, 811–816 (2006).

34. Meers, M. P., Janssens, D. H. & Henikoff, S. Pioneer Factor-Nucleosome Binding Events during Differentiation Are Motif Encoded. Mol. Cell 75, 562–575.e5 (2019).

35. Baranello, L., Kouzine, F., Sanford, S. & Levens, D. ChIP bias as a function of cross-linking time. Chromosom. Res. 24, 175–181 (2016).

36. Ernst, J. & Kellis, M. ChromHMM: automating chromatin-state discovery and characterization. Nat. Methods 9, 215–216 (2012).

37. Ernst, J. & Kellis, M. Chromatin-state discovery and genome annotation with ChromHMM. Nat Protoc 12, 2478–2492 (2017).

38. Uckelmann, M. & Davidovich, C. Not just a writer: PRC2 as a chromatin reader. Biochem. Soc. Trans. 49, 1159–1170 (2021).

39. Laugesen, A., Hojfeldt, J. W. & Helin, K. Molecular Mechanisms Directing PRC2 Recruitment and H3K27 Methylation. Mol Cell 74, 8–18 (2019).

40. Seneviratne, J., Ho, W. W., Glancy, E. & Eckersley-Maslin, M. A. A low-input high resolution sequential chromatin immunoprecipitation method captures genome-wide dynamics of bivalent chromatin. Epigenetics Chromatin 17, 3 (2024).

41. Suzuki, A. et al. DBTSS as an integrative platform for transcriptome, epigenome and genome sequence variation data. Nucleic Acids Res. 43, D87–D91 (2015).

42. Westermann, L. et al. Wildtype heterogeneity contributes to clonal variability in genome edited cells. Sci Rep-uk 12, 18211 (2022).

43. Bernhard, P. et al. Proteome alterations during clonal isolation of established human pancreatic cancer cell lines. Cell Mol Life Sci 79, 561 (2022).

44. Glancy, E., Ciferri, C. & Bracken, A. P. Structural basis for PRC2 engagement with chromatin. Curr. Opin. Struct. Biol. 67, 135–144 (2021).

45. Lasko, L. M. et al. Discovery of a selective catalytic p300/CBP inhibitor that targets lineage-specific tumours. Nature 550, 128–132 (2017).

46. Wang, J. et al. SPOP suppresses testicular germ cell tumors progression through ubiquitination and degradation of DPPA2. Biochem. Biophys. Res. Commun. 557, 55– 61 (2021).

47. Gupta, N. et al. A genome-wide screen reveals new regulators of the 2-cell-like cell state. Nat. Struct. Mol. Biol. 1–14 (2023) doi:10.1038/s41594-023-01038-z.

48. Iaco, A. D., Coudray, A., Duc, J. & Trono, D. DPPA2 and DPPA4 are necessary to establish a 2C-like state in mouse embryonic stem cells. Embo Rep 20, e47382–e47382 (2019).

49. Healy, E. et al. PRC2.1 and PRC2.2 Synergize to Coordinate H3K27 Trimethylation. Mol. Cell 76, 437–452.e6 (2019).

50. Blackledge, N. P. et al. PRC1 Catalytic Activity Is Central to Polycomb System Function. Mol. Cell 77, 857–874.e9 (2020).

51. Fursova, N. A. et al. Synergy between Variant PRC1 Complexes Defines Polycomb-Mediated Gene Repression. Mol. Cell 74, 1020–1036.e8 (2019).

52. Tamburri, S. et al. Histone H2AK119 Mono-Ubiquitination Is Essential for Polycomb-Mediated Transcriptional Repression. Mol. Cell 77, 840–856.e5 (2020).

53. Kalb, R. et al. Histone H2A monoubiquitination promotes histone H3 methylation in Polycomb repression. Nat. Struct. Mol. Biol. 21, 569–571 (2014).

54. Blackledge, N. P. et al. Variant PRC1 Complex-Dependent H2A Ubiquitylation Drives PRC2 Recruitment and Polycomb Domain Formation. Cell 157, 1445–1459 (2014).

55. Ohtomo, H. et al. H2A Ubiquitination Alters H3-tail Dynamics on Linker-DNA to Enhance H3K27 Methylation. J. Mol. Biol. 435, 167936 (2023).

56. Sugishita, H. et al. Variant PCGF1-PRC1 links PRC2 recruitment with differentiation-associated transcriptional inactivation at target genes. Nat. Commun. 12, 5341 (2021).

57. Kasinath, V. et al. JARID2 and AEBP2 regulate PRC2 in the presence of H2AK119ub1 and other histone modifications. Science 371, (2021).

58. Cooper, S. et al. Jarid2 binds mono-ubiquitylated H2A lysine 119 to mediate crosstalk between Polycomb complexes PRC1 and PRC2. Nat. Commun. 7, 13661 (2016).

59. Zhao, J. et al. H2AK119ub1 differentially fine-tunes gene expression by modulating canonical PRC1- and H1-dependent chromatin compaction. Mol. Cell 84, 1191–1205.e7 (2024).

60. Pasini, D. et al. Characterization of an antagonistic switch between histone H3 lysine 27 methylation and acetylation in the transcriptional regulation of Polycomb group target genes. Nucleic Acids Res. 38, 4958–4969 (2010).

61. Tie, F., Banerjee, R., Conrad, P. A., Scacheri, P. C. & Harte, P. J. Histone Demethylase UTX and Chromatin Remodeler BRM Bind Directly to CBP and Modulate Acetylation of Histone H3 Lysine 27. Mol. Cell. Biol. 32, 2323–2334 (2012).

62. Tie, F. et al. CBP-mediated acetylation of histone H3 lysine 27 antagonizes Drosophila Polycomb silencing. Development 136, 3131–3141 (2009).

63. Hogg, S. J. et al. Targeting histone acetylation dynamics and oncogenic transcription by catalytic P300/CBP inhibition. Mol. Cell 81, 2183–2200.e13 (2021).

64. Zhang, B. et al. A dynamic H3K27ac signature identifies VEGFA-stimulated endothelial enhancers and requires EP300 activity. Genome Res. 23, 917–927 (2013).

65. Zhan, X. et al. Glioma stem-like cells evade interferon suppression through MBD3/NuRD complex–mediated STAT1 downregulation. J. Exp. Med. 217, e20191340 (2020).

66. Hunt, G., Boija, A. & Mannervik, M. p300/CBP sustains Polycomb silencing by non-enzymatic functions. Mol. Cell 82, 3580–3597.e9 (2022).

67. Bernstein, B. E. et al. A bivalent chromatin structure marks key developmental genes in embryonic stem cells. Cell 125, 315–26 (2006).

68. Azuara, V. et al. Chromatin signatures of pluripotent cell lines. Nat Cell Biol 8, 532–8 (2006).

69. Glancy, E., Choy, N. & Eckersley-Maslin, M. A. Bivalent chromatin: a developmental balancing act tipped in cancer. Biochem. Soc. Trans. 52, 217–229 (2024).

70. Macrae, T. A., Fothergill-Robinson, J. & Ramalho-Santos, M. Regulation, functions and transmission of bivalent chromatin during mammalian development. Nat Rev Mol Cell Bio 1–21 (2022) doi:10.1038/s41580-022-00518-2.

71. Yan, P. et al. Genome-wide R-loop Landscapes during Cell Differentiation and Reprogramming. Cell Rep. 32, 107870 (2020).

72. Ikeda, H., Sone, M., Yamanaka, S. & Yamamoto, T. Structural and spatial chromatin features at developmental gene loci in human pluripotent stem cells. Nat. Commun. 8, 1616 (2017).

