## Supplemental Information for "Embryonic stem cell factors DPPA2/4 facilitate a unique chromatin state in non-small cell lung cancer"

#### Senevriatne et al. 2025

#### Extended Figure Legends

**Extended Figure 1, related to Figure 1**

**a)** GTEx normal adult tissue DPPA2/4 expression in boxplots (n=19616, units are in log10TPM+1).

**b)** DPPA2/4 expression in the TCGA TGCT cohort stratified by expression group (co-expressing and not co-expressing) (units are in log2RSEM+1 counts). Boxplots shown adjacent to the axes represent the expression of DPPA2 and DPPA4 in either group (median, whiskers represent the lower and upper 95% confidence intervals).

**c)** Pan-cancer molecular profile of DPPA2/4 across 31 cancer types from TCGA. The first panel summarises the proportion of patients with a copy number aberrations on the DPPA2/4 gene body (WES; copy number ≥ 3 gain, copy number ≤ 1 loss, copy number = 2 diploid). The second panel summarises the proportion of patients with somatic mutations on the DPPA2/4 gene body (WES). The third panel indicates the relative ploidy (copy number) of DPPA2/4 as boxplots for either DPPA2/4 (WES; log2RelativePloidy). The fourth panel indicates the gene expression of DPPA2/4 as boxplots for either DPPA2/4 (RNA-seq; log2RSEM). The fifth panel indicates the methylation level of DPPA2/4 gene promoters (TSS+/-1.5kb) as boxplots for either DPPA2/4 (methylation array; β-value). The sixth panel indicates the chromatin accessibility of DPPA2/4 gene promoters (DPPA2p1, DPPA4p1, DPPA4p2, DPPA4p3) or putative enhancers (DPPA4e1, DPPA4e2) as boxplots for either DPPA2/4 (ATAC-seq; log2Normalised Counts).

**d)** DPPA2/4 expression in the TCGA LGG, LUSC and LUAD cohorts stratified by expression group (co-expressing and not co-expressing) (units are in log2RSEM+1 counts). Boxplots shown adjacent to the axes represent the expression of DPPA2 and DPPA4 in either group (median, whiskers represent the lower and upper 95% confidence intervals).

**e)** Scatter plot of TCGA cancer cohort Cox proportional hazard modelling (CoxPH) of patient outcomes, where patients in each cohort was subdivided by the expression of either DPPA2 or DPPA4 as determined by RNA-seq (expression cut-off defined as the median of all DPPA2 or DPPA4 expressors (RSEM>0)). The x-axis and y-axis indicates the hazard ratio (HR) of CoxPH models (log10 transformed) for DPPA2 and DPPA4 respectively. Dots are coloured to indicate significant models (absolute HR > 1 and p-value < 0.05) for either/both DPPA2 and DPPA4 models, whiskers represent the lower and upper 95% confidence intervals of HRs for each model.

**f)** Select CoxPH models are shown for TCGA LGG, LUSC and LUAD cohorts when pateints were subdivided by DPPA2, DPPA4 or DPPA2+4 expression. The x-axis indicates the hazard ratio (HR) of CoxPH models (log10 transformed) with p-values of the model reported beside and each point highlighted by significance (p-value < 0.05).

**g)** Boxplots of DPPA2/4 promoter (TSS+/-1500bp) DNA methylation in a range of human cell line models spanning human development from the epigenetic roadmap consortium (ERM). DNA methylation is presented as fraction methylated, with groups of cell lines ordered by decreasing average DPPA2/4 promoter DNA methylation.

**h)** Dot plot visualising the correlation coefficient (biserial correlation for mutations and pearson correlation for all others) of DPPA2/4 gene expression (RNA-seq) correlated with their gene expression (RNA-seq), promoter DNA methylation (methylation arrays), gene copy number (WES), gene somatic mutation (WES) and chromatin accessibility at promoters or distal enhancers (ATAC-seq) in TCGA TGCT, LGG, LUSC and LUAD cancer cohorts. Significant correlations (BH-adjusted p-value < 0.05) are marked with an asterisk*. Those correlations for which there was no accessibility data or lack of mutations are marked with a question mark (?).

**i)** Kaplan-Meier survival curves of Balb/c nude mice xenografted with NCI-H1299 DPPA2/4 double overexpression lines (iFlag-DPPA2 and iV5-DPPA4). Lines were pre-treated with a vehicle control ($H_20$) or doxycycline 2µg/mL doxycycline (DOX) for 72h prior to engraftment and then xenografted mice were respectively kept on control or doxycycline feeds to maintain overexpression. Endpoint is tumour volume reaching 1000$mm^3$. P-value is computed from a log-rank test.
**j)** Tumour volumes ($mm^3$) of xenografted mice in **i**).
**k)** Volcano plot illustrating MSigDB C2 gene sets after a GSEA using all genes ranked by log2FC comparing DPPA2/4 co-expressing vs other NSCLC (LUAD+LUSC) tumours. Depleted gene sets are determined as those with NES < -1 and adjusted p-value < 0.05 and enriched gene sets are determined as NES > 1 and adjusted p-value < 0.05. Enriched gene sets falling into broad categories are annotated.

**Extended Figure 2, relating to Figure 2.**
**a)** Scatter plot of DPPA2 (x-axis) and DPPA4 (y-axis) gene expression across cancer cell lines (n=1393) in the CCLE (DepMap; RNA-seq) (units are log2TPM+1). Cell lines are coloured by whether DPPA2 and/or DPPA4 are expressed, where co-expressing cell lines are labelled.
**b)** Scatter plot of DPPA2 (x-axis) and DPPA4 (y-axis) protein levels across cancer cell lines (n=949) from Goncalves et al 2022 (Mass Spectrometry) (units are log2 Protein Intensity +1). Cell lines are coloured by whether DPPA2 and/or DPPA4 are detected, where co-detected cell lines are labelled.
**c)** Column graphs of relative DPPA2 and DPPA4 gene expression (relative to parental line and normalised to GAPDH & RPL19 housekeeping controls) in parental and isogenic NCI-H661 clones following CRISPR-Cas9 knockout clones and RT-qPCR (n=3 independent passages).
**d)** Violin plots of quantified median fluorescence intensity from DPPA2 and DPPA4 immunoflourescence for each isogenic cell line (2 clones per genotype, X images, N fields per image, X magnification).
**e)** Brightfield image of western blot membrane following reversible staining with the Ponceau S protein stain. The molecular ladders are present in the first and last lane. Six lanes indicate the 25% of the whole cell lysate inputs (Replicate 1-6) used for the immunoprecipitations. The next four lanes indicate a representative immunoprecipitation (Replicate 1 only eluted and blotted for, Replicate 2-6 were submitted for mass spectrometry following on-bead tryptic digests) with IgG (mouse), DPPA4 (rabbit), DPPA2 (mouse) and IgG (rabbit) antibodies respectively.
**f)** Western blots of above membrane following incubation with anti-DPPA2 or anti-DPPA4 antibodies. The expected sizes of DPPA2 and DPPA4 are annotated, as well as light (25 kDa) and heavy (55 kDa) chain IgG antibody fragments following elution.
**g)** Principal component analysis (PCA) of mass spectrometry data using LFQ-AUC values for all proteins (n=4 biological replicates/independent lysates per condition). Samples are coloured by IP type, with PC1 and PC2 representing the most variable components (28.85% and 15.63% respectively).
**h)** Scatter plots of detected nuclear proteins in the IP-MS data (subset of all proteins with an annotated nuclear localisation within the human protein atlas). Significantly enriched proteins (relative to IgG controls) are annotated (log2FC ≥ 1, BH-adjusted p-value < 0.05).
**i**) Representative histograms of mass photometry data of recombinant DPPA2-Myc/Flag and/or His-DPPA4 proteins, where the x-axis represents mass (kDa) and y-axis represents photometry counts. Called photometry peaks are annotated with median size (kDa), standard error (σ) and skewness. Schematic of DPPA2 (red) and DPPA4 (purple) oligomeric complexes are shown. The experiments were repeated three times independently.
**j)** Bar plots summarising mass photometry peak sizes of recombinant DPPA2-Myc/Flag and/or His-DPPA4 proteins (n=three to four experimental replicates) at different complex stoichiometries

**k)** Venn diagrams summarising the overlap of our IP-MS findings with other studies; For DPPA2-IP: 1) Eckersley-Maslin et al. 2020 NSMB, overexpression of DPPA2/4-GFP, IP using GFP-trap followed by RIME-MS in mESC's, 2) Hernandez et al. 2018 Cell Stem Cell, overexpression of 3xFlag-DPPA2, following Flag-IP and MS in mESCs. For DPPA4-IP: 1) Eckersley-Maslin et al. 2020 NSMB as above, 2) Oliviera et al. 2014, endogenous DPPA4-IP followed by MS in NT2 human cells.
**l,m,n,o)** Summary of enrichment (log2FC IP vs IgG) in IP-MS for detected proteins belonging to PRC1, PRC2, COMPASS or histones. Significant enrichments are denoted by an asterisk (* BH adjusted p-value < 0.05 and log2FC ≥ 1).
**p)** Enrichment (log2FC IP vs IgG) of H2A peptides in IP-MS
**q)** Line curves summarising the binding of DPPA2-Myc/Flag and/or His-DPPA4 proteins to nucleosomes in EMSA experiments (n=3 experimental replicates), represented as the percentage of total nucleosomes bound. Error bars represent the standard error of the mean.

**Extended Figure 3, relating to Figure 3**
**a)** Heatmaps of genomic regions centred on DPPA2/4 peaks as determined by ChIP or CUT&Run (resized to +/-5kb from the centre of each peak, split into 100 equal windows) as well as a set of background regions not bound by DPPA2+4 for comparison (DPPA2 (n=12866), DPPA4 (n=5773), DPPA2+4 (n=16620) and background (n=1000). Columns represent the double crosslinked input ChIP control (n=1), averaged mouse/rabbit IgG CUT&Run control (n=1 each), average DPPA2 (n=2) ChIP (MAB4356), DPPA2 (n=1) CUT&Run (MAB4356), average DPPA4 (n=2) ChIP (AP1438a), DPPA4 (n=1) CUT&Run (AP1438a), average DPPA4 (n=2) ChIP (ab154642) , DPPA4 (n=1) CUT&Run (ab154642), DPPA4 (n=1) CUT&Run (AP21202c). ChIP and CUT&Run CPM/bp values were scaled per bin across all regions for visualisation.
**b)** Venn diagram of ChIP (overlapping peaks in 2 replicates, determined as those peaks overlapping by atleast 25%) and CUT&Run (1 replicate) peak overlaps for DPPA2 (MAB4356) and DPPA4 (AP1438a) using MACS2 (FDR < 0.05 and log2FC > 1 compared to matched Input or IgG controls).
**c)** Genome-wide pairwise Pearson correlations of ChIP-seq and CUT&Run (DPPA2 (n=2) ChIP (MAB4356), DPPA2 (n=1) CUT&Run (MAB4356), DPPA4 (n=2) ChIP (AP1438a), DPPA4 (n=1) CUT&Run (AP1438a), DPPA4 (n=2) ChIP (ab154642), DPPA4 (n=1) CUT&Run (ab154642), DPPA4 (n=1) CUT&Run (AP21202c) fold enrichment over the matched input or IgG control) partitioned into 200bp bins genome-wide. Dots are coloured and sized by Pearson correlation coefficients (r) and are ordered by hierarchical clustering with red squares denoting major clusters.
**d)** Top 5 HOMER *de novo* motifs from DPPA2+4 ChIP/CUT&Run peaks (n=16620) relative to a size and gc-content matched set of background genomic regions (n=16620). All regions were resized to 1000bp prior to analyses.
**e)** HOMER genomic annotations of DPPA2 and DPPA4 ChIP/CUT&Run peaks (determined by MACS2, FDR < 0.05, log2FC > 1). Promoter-TSS is defined as TSS+/-1500bp. The numbers atop each column indicates the total number of DPPA2/4 peaks (for ChIPs these are the number of overlapping peaks (>25%) between two replicates, for CUT&Run these are from one replicate each).
**f)** Gene ontology (GO-Biological Processes) enrichment analysis of DPPA2+4 bound gene promoters as determined by clusterProfileR. The red line indicates the significance threshold at adjusted p-value 0.05.
**g)** HOMER genomic annotations of H3K4me3, H3K27ac, H3K4me1, H2AK119ub, H3K27me3, H3K9me3, H3K36me2 and H3K36me3 ChIP peaks (determined by SICER2, FDR < 0.05, log2FC > 1). Promoter-TSS is defined as TSS+/-1500bp. The numbers atop each column indicates the total number of peaks (these are the number of overlapping peaks (>25%) between two replicates).

**h)** Genomic track plot of the chr 5:93429332-93810092 (hg38) locus, where a gene annotation track is provided at the top, followed by CpG island annotations, the double crosslinked input ChIP control (n=1), DPPA2 (n=2), DPPA4 (n=2) ChIP, single crosslinked input ChIP control (n=1), H3K4me3 (n=2), H3K27ac (n=2), H3K4me1 (n=2), H2AK119ub (n=2), H3K27me3 (n=2), H3K36me3 (n=2), H3K36me2 (n=2), H3K9me3 (n=2) ChIP and DNA methylation (WGBS) (n=3), followed by consensus DPPA2+4 peak regions annotated at the bottom. Scales are in CPM/bp for ChIPs and percentage methylated DNA for WGBS (0-100%).
**i)** 17-state chromatin model as determined by ChromHMM modelling. The first panel presents emission probabilities that summarise the association of each mark with each chromatin state. The second to fourth panel indicates the scaled overlap of each chromatin state with annotated features in the genome, note that RefSeq annotations are split according to expressed and repressed genes in NCI-H661, as determined by RNA-seq. The fifth panel indicates transition probabilities that indicate the relatedness between states. The sixth panel represents the scaled enrichment of each signal flanking the TSS+/-2kb.
**j)** H3K4me3-H2AK119ub reChIP peak calling and classification workflow. Here we overlapped peaks from 2 replicates (>25% overlap), and then for reChIP's overlapped consensus H3K4me3-H2AK119ub and H2AK119ub-H3K4me3 peaks to arrive at a final set of n=4663 reChIP peaks. These peaks were further subclassified using single H3K4me3 or H2AK119ub peak sets to classify peaks as either high confidence (HC, enriched in all conditions), H3K4me3-biased (K4b, enriched in H3K4me3, but not H2AK119ub), H2AK119ub-biased (K119b, enriched in H2AK119ub, but not H3K4me3) and low confidence (LC, not enriched in single ChIPs).

**Extended Figure 4, related to Figure 4**
**a)** PCA analysis of factorised chromatin states (ChromHMM) at regions not bound by DPPA2+4 (n=16620) in NCI-H661 cells across all NSCLC cell lines (n=27).
**b)** Schematic of doxycycline inducible shRNA pCLIIi system and RT-qPCR analyses of DPPA2 and DPPA4 in stable NCI-H661 shRNA cell lines following induction with a vehicle control ($H_2O$) or 2µg/mL doxycycline (DOX) for 7 days. The error bars represent the standard error of the mean of n=3 biological replicates.
**c)** RT-qPCR analyses of NCI-H661 cells following siRNA mediated DPPA2 or/and DPPA4 knockdown for 4 days. The error bars represent the standard error of the mean of n=3 biological replicates.
**d-f)** Scatter plots of differential RNA-seq (n=3 for all conditions) in NCI-H661 following 4 days of siRNA mediated DPPA2 or/and DPPA4 knockdown. Differentially expressed genes are those with an FDR < 0.05 and absolute log2FC of ≥ 1 as determined by edgeR-TMM analyses between DPPA2 and/or DPPA4 siRNA vs control siRNA. The y-axes indicate the log2FC for siDPPA2/4 vs siControl, and the x-axes are the counts per million (logCPM) in the siControl for all genes.
**g-i)** Scatter plots of differential whole genome bisulfite-seq (WGBS, n=3 for all conditions) in NCI-H661 following 4 days of siRNA mediated DPPA2 or/and DPPA4 knockdown. Differentially methylated CpGs are those with an FDR < 0.05 as determined by edgeR analyses between DPPA2 and/or DPPA4 siRNA vs control siRNA. The y and x-axes indicate the fraction of methylated reads for a given CpG in siDPPA2/4 vs siControl respectively. Note that only variable CpG's with coverage (i.e. those CpG's that weren't solely hypo(0)/hypermethylated(1) across all samples and those CpG's with atleast 1 read across all samples) were tested.
**j)** Heatmaps of genomic regions centred on H2AK119ub differential peaks (lost) in shREN (non-targeting control) and shDPPA2 (resized to +/-5kb from the centre of each peak, split into 100 equal windows) as well as a set of H2AK119ub unchanged regions for comparison (depleted in both shDPPA4 and shDKD (n=1070), shDPPA4 only (n=2918), shDKD only (n=859) and unchanged (n=500). Columns represent average DPPA2 (n=2) and DPPA4 (n=2) ChIP, DPPA2 (n=1) and DPPA4 (n=1) CUT&Run and averaged ATAC-seq, H3K4me3, H3K27me3 and H2AK119ub ChIP-seq for shREN and shDPPA2 (n=2 each for $H_2O$ and DOX). DPPA2/4 ChIP and

CUT&Run CPM/bp values were scaled per bin across all regions for visualisation, whereas shREN and shDPPA2 ATAC-seq and ChIP-seq represent unscaled CPM/bp.
**k)** Proportional bar plot of HOMER genomic annotations in all H2AK119ub regions or in subsets of these regions that were depleted in shDPPA4/shDKD. The total number of peaks per category are annotated atop each bar.
**l)** Heatmap of known motifs enriched in shDPPA4/shDKD H2AK119ub depleted regions as determined by monaLisa analyses. Depleted regions were first resized to 4000bp, corresponding to the approximate median length of all depleted domains. Size and gc-matched regions were then sampled from the genome as background regions for the analysis (bck, n=859). The heatmap presents all motifs that were enriched over background (log2FC > 0.25 and BH-adjusted p-value < 0.05) in any depleted category. Motifs as well as their gc content are provided to the left of the plot.
**m)** Gene ontology (GO-Biological Processes) enrichment analysis of gene promoters overlapping H2AK119ub depleted regions (TSS+/-1500bp) as determined by clusterProfileR. Coloured dots represent terms passing the significance threshold at adjusted p-value 0.05 and their size reflects the proportion of genes represented by each depleted category and the total number of genes in the geneset.

**Extended Figure 5, related to Figure 5**
**a)** Venn diagram of H3K27ac (n=2), H3K27me3 (n=2) and SUZ12 (n=1) ChIP peak overlaps (overlapping peaks in 2 replicates for H3K27ac and H3K27me3, determined as those peaks overlapping by at least 25%).
**b)** Expression of CDH6 in control (grey) vs A-485 (blue) treated cells
**c-e)** genome browser views (left) and expression (right) for DPPA2/4-dependent H2AK119Ub class I (GPC4, panel c) and class II (MN1, panel d; PCDH7, panel e) promoters

### Extended Materials & Methods

#### Cell culture

Non-small cell lung cancer cell lines NCI-H661 and NCI-H1299 were cultured in RPMI-1640 (Gibco) growth medium supplemented with 10% heat-inactivated fetal bovine serum (FBS) and 1x penicillin/streptomycin at 37℃, 5% $CO_2$. E14 mouse embryonic stem cells were cultured on feeder-free gelatinised plates at 37 ℃, 5% $CO_2$ using standard serum/LIF conditions (high-glucose DMEM supplemented with 15% FBS, 1x GlutaMax, 1x penicillin/streptomycin, 0.1mM nonessential amino acids, 50mM beta-mercaptoethanol and LIF (made in house in HEK293 cells and titrated for optimal ESC growth)). Cells were routinely passaged to maintain exponential growth, using Trypsin-Low EDTA (LE) Express (Gibco) to detach cells each time. Cells were regularly tested for mycoplasma contamination using the Mycoplasma PCR Detection Kit (abcam, #ab289834). NSCLC cell lines were also STR profiled to ensure authenticity. NCI-H661 and NCI-H1299 were purchased from ATCC, and E14 mouse embryonic stem cells were a gift from Wolf Reik's laboratory.

#### Inducible shRNA and cDNA construct generation and stable cell line generation

For inducible knockdown experiments, miR30 based shRNAs were expressed using a tetracycline-inducible *PiggyBac* transposon system (pCLIIPi-iRFP-shRNA-pGK-mVenus), as previously described[1] (Eckersley-Maslin *et al*. 2020). Established shRNA sequences[2] against either; Renilla luciferase (non-targeting control) (#713), human DPPA2 (#760) or DPPA4 (#1993) were cloned by first annealing complementary oligo's and ligating them into a digested pMSCV-miR30 backbone[3] , followed by PCR, digestion and ligation into the pCLIIPi backbone. Stable cell lines were generated by transfecting NCI-H661 cells with the shRNA vectors along with *PiggyBac* transposase (pRM1024)[4] , waiting for 2 weeks, then sorting mVenus positive cells twice with expansion in between, collecting the top 50% of mVenus positive cells each time as a bulk population for further experiments. The mVenus DPPA4 shRNA construct underwent additional cloning to replace mVenus with tagBFP to allow for selection of cells containing both DPPA2 and DPPA4 inducible shRNA. To generate the double inducible knockdown line, stable shDPPA2 (mVenus+) cells were transfected with the shDPPA4 (tagBFP) vector and selected as above for both mVenus+/tagBFP+ cells. For inducible overexpression experiments the above vector backbones were cloned to replace the (iRFP-shRNA) fragment downstream of the tet-response element (TRE) with cDNA encoding human DPPA2 (ORF:NM_138815) or DPPA4 (ORF:NM_018189) with additional N-terminal Flag or V5 tags respectively (gene fragments synthesised by IDT). Stable cell lines were generated in the same way as above for shRNA lines for the NCI-H1299 cell line (which does not express DPPA2/4). All cell sorting was undertaken on FACSAria Fusion flow cytometry instruments (BD). To induce either shRNA or cDNA expression, cells were treated with 2μg/ml doxycycline (Dox) for 3-7 days and knockdown or overexpression were confirmed by RT-qPCR or western blot analysis. All vectors were sequence verified by either Sanger (shRNA constructs) or Plasmidsaurus (cDNA constructs) sequencing. shRNA sequences are in Table 1 below.

#### Generation of CRISPR-Cas9 knockout cell lines

NCI-H661 cells were electroporated using the Neon™ Transfection System (ThermoFisher) with Cas9 RNP (Horizon, CAS12205) loaded with a single guide RNAs targeted to DPPA2 (Exon 6: 5'-GCGATGTTCGAGGAAACGCA-3') or DPPA4 (Exon 3: 5'-CGGTGAATCAGATTAACAGG-3') respectively. Post electroporation, cells were seeded at low density (1 cell/well in a 384-well plate) to isolate monoclonal populations. After 3 weeks in single-cell dilution, expanded populations were screened by PCR following gDNA extraction and edits identified by Sanger sequencing. Clones with an indel knockout on all alleles were expanded. To generate DPPA2/4 double knockout clones DPPA2 single KO clone c40 was subject to DPPA4 CRISPR-Cas9 KO as above, and then subcloned and genotyped as above. The generation and validation of these isogenic cell lines were undertaken by Horizon (CLPP1690). Knockouts were further validated using western blot analyses.

**siRNA transfection**

siRNA knockdowns for DPPA2, DPPA4, both DPPA2+4 were performed using 10uL Lipofectamine 2000 (ThermoFisher, #11668027) and 200pmol of non-targeted and DPPA2/4 targeted siRNA twice over 96 hours (first dose at day 0, second dose at day 2, collection on day 4) using SMARTpool siRNA (containing 4 siRNAs per pool) (Horizon/Dharmacon, Non-targeting; #D-001810-10-05, DPPA2; #L-018977-01-0005, DPPA4; #L-020766-01-0005). Lipofectamine 2000 and siRNA complexes were first formed with reduced-serum OptiMEM medium (Gibco) for 20 mins at room temperature, prior to addition to cells grown in complete medium. Knockdown efficiency was validated by RT-qPCR following RNA extraction. siRNA sequences are in Table 2 below.

**gDNA extraction and Whole Genome Bisulfite Library preparation**

Following treatment or passaging, NCI-H661 cells were harvested, washed once with PBS, stored in RLT+ lysis buffer (Qiagen) and snap frozen. gDNA was extracted from cells using an AllPrep RNA/DNA extraction kit (Qiagen #80204) according to the manufacturer's instructions. DNA libraries were generated using the NEBNext End Prep kit (NEB E7370, E7535) and bisulfite converted using the EZ DNA Methylation Gold Kit (Zymo Research #D5005, D5006) according to the manufacturer's instructions. Libraries were amplified using KAPA HiFi Uracil+ (KK2801/2) using NEBNext Universal and Index primers, cleaned up using 0.8x Ampure beads and pooled for sequencing.

**RNA extraction, cDNA synthesis and RNA-seq library preparation**

Following treatment or passaging, cells were enzymatically detached, washed once with PBS and snap frozen as a pellet or stored in RLT+ lysis buffer. RNA extraction was performed on cell pellets (NEB) or lysates (Qiagen), using either the Monarch Total RNA miniprep kit (NEB #T2010S) or AllPrep RNA/DNA extraction kit (Qiagen #80204) with DNAse I treatment according to the manufacturer's instructions. 0.5-1µg of RNA was used for cDNA synthesis via reverse transcription using the LunaScript RT kit (NEB #E3010). RNA was quantified using the Qubit™ RNA High Sensitivity (HS) Assay Kit (Invitrogen) on a Qubit 3.0 fluorometer. Following RNA quantitation, 05-1µg of RNA was used as input for PolyA+ directional RNA-seq library preparation using the NEBNext Ultra II Directional RNA-seq Kit (#E7765, NEB) with the PolyA mRNA magnetic isolation module (#E7490, NEB) according to manufacturer instructions.

**Whole cell protein extraction, quantification and immunoprecipitation**

Snap frozen NCI-H661 cell pellets ($5x10^6$ cells) were lysed in ice-cold RIPA lysis buffer (150mM NaCl, 50mM Tris-HCl, 1% IGEPAL CA-630 (Sigma), 0.5% sodium deoxycholate, 0.1% SDS in $H_2O$) for 30 minutes on ice with or without sonication for 30mins. Following centrifugation at 11,000xg to pellet debris, protein from whole cell lysates (WCL) were quantified using the BioRad protein quantitation assay (Bio-Rad, #5000006) and measured on a Cytation 3 (BioTek) plate reader using known BSA serial dilutions as a standard curve at 595nm (colorimetric). 430µg of protein WCL were used for each set of immunoprecipitations and were diluted to 165uL in PBST (PBS + 0.01% Tween-20). 100µL of Protein-G Dynabeads (Thermo) were washed with Citrate Phosphate Buffer (25mM citric acid, 50mM dibasic sodium phosphate dihydrate in $H_2O$ calibrated to pH5.0) twice followed by resuspension in 260uL of PBST. Diluted WCL were then pre-cleared to remove proteins that non-specifically bind to Protein-G Dynabeads by adding 55uL of washed beads to the diluted WCL and incubating them at 4C for 1hr on a rotator, followed by transfer of the pre-cleared eluate to a new tube. 2µg of antibodies (DPPA2 (Mouse); Sigma #MAB4356, DPPA4 (Rabbit); abcam #ab154642) or IgG's isotype controls (Mouse; Invitrogen #31903, Rabbit; Invitrogen #02-6102) diluted to 150uL in PBST were conjugated to 50uL of washed beads by incubating them together at 4C for 1hr on a rotator. Antibody-bead conjugates were washed two times with PBST and then resuspended in 55uL PBST. Immunoprecipitations were then performed by incubating 50uL of antibody-bead conjugates with 50uL of pre-cleared WCL overnight at 4C whilst rotating, the rest of the pre-cleared WCL was kept as an input control. Following immunoprecipitation, beads were washed five times with PBS on ice, and beads were snap frozen on dry ice as a pellet for eventual on-bead tryptic digest for mass spectrometry or eluted in NuPAGE LDS sample buffer (Invitrogen, #NP0007) at 70C for 10 minutes for western blot analysis.

**Mass spectrometry and analysis**

Enriched proteins were digested off the beads overnight at 37C in 50uL of 2M urea containing 1mM tris(2-carboxyethyl)phosphine and 4mM 2-chloroacetamide with 0.4ug of sequencing grade trypsin (Promega, #V5280) and LysC (Wako, #125-05061). The peptides were purified using SDB-RPS microcolumns, washed with 99% isopropanol containing 1% trifluoracetic acid (TFA) followed by 5% acetonitrile containing 1% TFA and the eluted with 80% acetonitrile containing 1% ammonium hydroxide. Peptides were dried by vacuum centrifugation and resuspended in 2% acetonitrile containing 0.1% TFA. Peptides were separated on a Dionex 3500 nanoUHPLC, coupled to an Orbitrap Lumos mass spectrometer via electrospray ionization in positive mode with 1.9 kV at 275 °C and RF set to 30%. Separation is achieved on a 50 cm × 75 µm column packed with C18AQ (1.9 µm) over 40 min at a flow rate of 300 nL/min. Peptides were eluted over a linear gradient of 3–40% Buffer B (Buffer A: 0.1% v/v formic acid; Buffer B: 80% v/v acetonitrile, 0.1% v/v FA) and the column was maintained at 50°C. The instrument was operated in data-independent acquisition (DIA) mode, with an MS1 spectrum acquired over the mass range 350–1,400 m/z (60,000 resolution, 100% automatic gain control (AGC), and 45 ms maximum injection time) followed by sequential MS/MS spectra across 13.7 m/z isolation windows with 1 m/z overlap covering the full mass range. MS/MS data will be acquired with higher-energy collisional dissociation (HCD) fragmentation (15,000 resolution, 2000% AGC, 55 ms maximum injection time, and normalized collision energy 30%). Data were processed in Spectronaut 1 (v7.6.230428.55965) with default setting against the Homo sapien protein FASTA sequences in the Uniprot database and filtered to 1% FDR at the PSM, peptide and protein level.

Raw label-free quantitation values (LFQ-AUC) values were subject to down-shifted imputation, whereby missing values for a given protein were imputed with a low intensity LFQ-AUC value based on the normalised distribution of the dataset. Following imputation, LFQ-AUC values were centered for each protein by log2 transformation and subtraction of the median across all samples. To determine enrichment statistics, unpaired Student's t-tests were performed between the IP conditions (DPPA2, DPPA4) and matched control IgG (Ms, Rb) respectively, followed by p-value adjustment to account for multiple comparisons using the Benjamini-Hochberg method.

**Subcellular fractionation**

Freshly harvested NCI-H661 shRNA cells following 7 days of induction ($2x10^6$ cells) were washed with ice-cold PBS once. Cells were pelleted at 400xg and washed three times in 200uL cytoplasmic extraction buffer (50mM HEPES, 140mM NaCl, 1mM EDTA, 10% Glycerol, 0.1% IGEPAL CA-630, 0.25% Triton-X, 1mM DTT in $H_2O$) on ice, with the first wash being kept as the cytoplasmic fraction after 10 mins of incubation. Nuclei were then washed three times in 80uL of nuclear extraction buffer (10mM Tris-HCl, 200mM NaCl, 1mM EDTA, 0.5mM EGTA in $H_2O$) on ice to release soluble nuclear proteins, of which the first wash was kept as the nuclear (soluble) fraction after 10 mins of incubation. Insoluble chromatin was then resuspended in 200uL chromatin extraction buffer (50mM Tris-HCl, 20mM NaCl, 1mM $MgCl_2$, 0.1% SDS, 1:1000 benzonase (Merck #E1014)) on ice and rotated at room temperature for 20 mins, followed by a spin at 16,000xg at 4C to pellet debris, of which the supernatant was kept as the chromatin (insoluble) fraction. All extraction buffers were freshly supplemented with 1x protease inhibitor tablet (). All intermediate spins between the first and last incubation were at 11,000xg at 4C. Successful cytoplasm-chromatin fractionation was confirmed by performing gel electrophoresis, transfer to nitrocellulose membranes, staining with ponceau S and observing the presence/absence of histones (H3, H2A/B, H4) between 10-25 kDa in each fraction.

**Gel electrophoresis, transfer and western blots**

Following protein extraction and protein quantitation (as above), 50ug of protein were prepared with NuPAGE 4xLDS sample buffer (Invitrogen) and subsequently denatured at 85C for 5 mins. Samples were loaded into 10,12 or 17-well NuPAGE 4-12% gradient Bis-Tris gels (Invitrogen) placed within an XCell SureLock electrophoresis cell with either NuPAGE MES or MOPS running buffer (Invitrogen). 5uL of PageRuler plus prestained protein ladder (ThermoFisher) was included in a well for each blot. Gel electrophoresis was performed at 90V for 5 mins, followed by 120V for 40-50 mins at room temperature. Proteins were then transferred from the gel to a nitrocellulose membrane using the XCell II Blot Module (Invitrogen) via wet transfer in NuPAGE transfer buffer (Invitrogen) containing 10% methanol for 1hr 30V at room temperature. To confirm transfer of proteins membranes were stained with Ponceau S (Merck). Membranes were then blocked for 2-4 hr at room temperature in 5% w/v Bovine Serum Albumin (A4737) or 0.1% skim-milk w/v in TBST (20mM Tris-HCl, 150mM NaCl, 0.1% Tween-20 in H2O), followed by a single wash in TBST for 5 mins. Membranes were divided for probing of multiple proteins at distinct molecular weights. Membranes were incubated with primary antibody at varying dilutions (1/2000 Flag (#F1804, Merck), 1/5000 V5 (#R960-25, Invitrogen), 1/1000 DPPA2 (Merck, #MAB4356), 1/1000 DPPA4 (abcepta, #AP1438a, abcepta, #AP21202c, abcam, #ab154642), 1/10000 H3K4me3 (CST,#9751), 1/10000 H3K27me3 (CST, #9733), 1/10000 H2AK119ub (CST, #8240), 1/10000 H3K27ac (abcam, #AB4729), 1/10000 total H2A (CST, #12349), 1/10000 total H3 (CST#, 9715), 1/5000 Vinculin (CST, #13901) in 0.1% skim-milk w/v in TBST overnight at 4C whilst rotating. Membranes were then washed three times in TBST for 5 mins each time, followed by incubation with HRP conjugated anti-rabbit (abcam #ab205718) or anti-mouse (abcam #ab205719) secondary antibodies at a 1:2000 dilution in 5% w/v Bovine Serum Albumin (A4737) or 0.1% skim-milk w/v in TBST for 2-4 hrs at room temperature whilst rocking. Membranes were washed three times with TBST for 5mins each, then incubated in Clarity chemiluminescent ECL substrate (Bio-Rad) for 2 mins followed by chemiluminescence visualisation on the iBright 1500 instrument (Invitrogen). When required, membranes were stripped of bound antibodies by incubation with Restore western blot stripping buffer (ThermoFisher) for 30 mins at room temperature whilst rocking, followed by TBST washes, re-blocking and probing of antibodies as above.

**Immunofluorescence, image acquisition and analysis**

NCI-H661 cell lines were seeded on sterilised glass coverslips placed within 6-well plates. When cells were ~70% confluent medium was aspirated, wells were washed twice with PBS, fixed in 4% formaldehyde in PBS for 10-15 mins at room temperature. Cells were washed twice with PBS and then permeabilised with 0.1% Triton-X in PBS for 5-10 minutes at room temperature. Cells were washed twice with PBS and then blocked with 3% BSA in PBS for at least 3 hours at room temperature whilst rocking. Primary antibodies (1/50 DPPA2 (Merck, #MAB4356), 1/50 DPPA4 (abcam, #ab154642)) were diluted in 3% BSA in PBS and coverslips were placed downward between parafilm for uniform staining 60-90 minutes at room temperature in a humidity chamber. Coverslips were washed 3x in PBS followed by incubation in fluorophore conjugated secondary antibodies anti-rabbit Alexa 594 (ThermoFisher, #A11037) or anti-mouse Alexa 488 (ThermoFisher #A32723) at 1/1000 in 3% BSA in PBS for 1hr at room temperature. Following 3 washes in PBS slides were counterstained with DAPI (ThermoFisher, #62248) at 1/10,000 in PBS for 1min at room temperature. Slides were mounted onto SuperFrost microscope slides with SlowFade Gold antifade (Invitrogen) and sealed with nail polish. Slides were imaged on an Olympus FV3000 confocal microscope. Image analysis was performed using ImageJ.

**Flow cytometry analysis and cell sorting**

Cells were resuspended in FACS buffer (2% FBS, 4mM EDTA in PBS) prior to flow cytometry analysis using the LSR II (BD) flow cytometer or sorted using the FACSAria Fusion flow cytometer (BD).

**Chromatin immunoprecipitation (ChIP)**

For DPPA2, DPPA4, KDM2A and SUZ12 ChIPs $4x10^6$ NCI-H661 cells were utilised per ChIP and underwent double-crosslinking prior to chromatin isolation. All other ChIPs against histone modifications (H3K4me3, H3K4me1, H3K27ac, H3K27me3, H2AK119ub, H3K9me3, H3K36me3, H3K36me2) utilised $5x10^5$ NCI-H661 cells per ChIP and underwent single-crosslinking. For double-crosslinking, cells were harvested, washed once with PBS, and resuspended in 10mL of PBS with 5mM EDTA on ice. A fresh stock of 0.24M disuccinimidyl glutarate (ThermoFisher #20593) was reconstituted in DMSO and added to the suspension of cells to reach a final concentration of 2mM, which were then incubated at room temperature whilst rocking for 45 mins to crosslink cells. Cells were pelleted at 3200xg for 5 mins and resuspended in 9.375mL of PBS with 5mM EDTA. Cells undergoing only single-crosslinking were resuspended in 9.375mL PBS at this point. 625uL of fresh 16% formaldehyde (methanol-free) (ThermoFisher, #28908) were added to each cell suspension to reach a final concentration of 1% and then incubated on a rocker for 12.5 mins at room temperature to crosslink cells. Crosslinkers were then quenched by supplementing cell suspensions with 1M ice-cold glycine to reach a final concentration of 125mM and rocked for a further 5 mins at room temperature. Crosslinked cells were then pelleted at 3200xg for 5mins at 4C, washed with 1mL ice-cold PBS with 5mM EDTA and protease inhibitors, and then immediately snap frozen with dry ice and stored at -80C until ChIPs were performed. For treatment-based ChIP experiments (following vehicle controls, DOX or A-485 treatment), 50,000 (10%) single-crosslinked E14 mouse ESCs were additionally spiked-in prior to each ChIP to act as exogenous normalisation references in case global changes to histone modifications were observed.

Crosslinked cell pellets were washed with 1mL ice-cold NP buffer (10mM Tris-HCl, 1M sorbitol, 50mM NaCl, 5mM $MgCl_2$, 1mM $CaCl_2$, 0.1% IGEPAL CA-630, 0.1% Tween-20 in $H_2O$ freshly supplemented with 0.385μM beta-mercaptoethanol, 1.82mM spermidine, 1x protease inhibitor tablet) twice to isolate nuclei, with nuclei spun at 2000xg for 5 mins at 4C each time for pelleting. Nuclei were resuspended in 900uL ChIP buffer (20mM Tris-HCl, 2mM EDTA, 150mM

NaCl, 0.6% SDS, 1% Triton-X in $H_2O$ freshly supplemented with 0.1µM PMSF, 1x protease inhibitor tablet), transferred to a milliTUBE AFA Snap Fibre sonication vial (Covaris, #520135) and left to incubate on ice for 1hr. Chromatin was then sheared to ~200bp fragments using the ME220 sonicator (Covaris) (4C, 75% power, 15% duty factor, 1000 cycles) for either 30 mins (double-crosslinked) or 20 mins (single-crosslinked). Debris were pelleted at 12,000xg for 15 mins at 4C, and the supernatant containing sheared chromatin was transferred to a new tube. Sheared chromatin was regularly checked through gel electrophoresis to ensure predominantly mononucleosomal fragments. For double-crosslinked ChIPs Protein-G dynabeads were used, and for single-crosslinked ChIPs Protein-A dynabeads were used. 50uL of beads were washed twice in 1mL of low-salt wash buffer (20mM Tris-HCl, 2mM EDTA, 150mM NaCl, 0.1% SDS, 1% Triton-X in $H_2O$) and resuspended in 100uL of low-salt buffer. To pre-clear chromatin, 20uL of washed beads were added to 180uL sheared chromatin and then incubated at 4C for 5 hours on a rotator, followed by transfer of the supernatant to a new tube. To conjugate antibodies to beads, 0.5-4µg of antibodies were diluted to 320uL with ChIP dilution buffer (100mM NaCl, 0.02% sodium azide, 100mM Tris-HCl, 5mM EDTA, 5% Triton-X in $H_2O$ supplemented with 1x protease inhibitor tablet) and were left to conjugate with 80uL of washed beads at 4C for 5 hours whilst rotating (IgG controls (0.5-4ug Mouse IgG (Invitrogen, #31903), 0.5-4ug Rabbit IgG (Invitrogen, #012-6102)) or antibodies (4ug DPPA2 (Sigma, #MAB4356), 4ug DPPA4 (abcepta, #AP1438a, abcepta, #AP21202c, abcam, #ab154642), 1ug SUZ12 (CST, #3737), 0.5ug H3K4me3 (CST, #9751), 0.5ug H3K27me3 (CST, #9733), 0.5ug H2AK119ub (CST, #8240), 0.5ug H3K4me1 (CST, #5326T), 1ug H3K27ac (abcam, #ab4729), 1ug H3K9me3 (abcam, #ab8898), 0.5ug H3K36me2 (abcam, #ab9049), 0.5ug H3K36me3 (CST, #4909)). Antibody-bead conjugates were washed with 1mL of ChIP dilution buffer and subsequently resuspended with 180uL of pre-cleared chromatin and left to incubate overnight whilst rotating at 4C, 9uL of the remaining pre-cleared chromatin was set aside as an input control. Beads were then washed three times with 200uL low-salt wash buffer, three times with 200uL high-salt wash buffer (20mM Tris-HCl, 2mM EDTA, 500mM NaCl, 0.1% SDS, 1% Triton-X in $H_2O$), two times with 200uL LiCl wash buffer (250mM LiCl, 1% sodium deoxycholate, 10mM Tris-HCl, 1mM EDTA, 1% IGEPAL CA-630) and once with 200uL 10mM Tris-HCl pH7.5. Beads as well as input controls were then resuspended/diluted in 50uL ChIP elution buffer (1% SDS, 0.1M $NaHCO_3$ in $H_2O$). Samples were subject to a RNAse A digest (5uL 20mg/mL NEB) for 1hr at 37C, then a Proteinase K digest (5uL 800U/mL NEB) for 1hr at 37C and crosslinks were reversed overnight by incubation at 65C on a thermomixer (1200RPM). DNA was then enriched from eluates using a 1.8X bead cleanup (SergiLabs) and eluted in 30uL $H_2O$ for downstream qPCR analyses and DNA library preparation.

**Sequential chromatin immunoprecipitation (reChIP)**

For each reChIP 1x10^6 single-crosslinked NCI-H661 cells were used. Pellets of single-crosslinked NCI-H661 cells were washed twice with 1mL NP buffer (10mM Tris-HCl, 1M sorbitol, 50mM NaCl, 5mM $MgCl_2$, 1mM $CaCl_2$, 0.1% IGEPAL CA-630, 0.1% Tween-20 in $H_2O$ freshly supplemented 1x protease inhibitor tablet) on ice. Nuclei were pooled and resuspended in 800µL ChIP buffer, then left to incubate on ice for at least 5 minutes. Chromatin was sheared (as above for single-crosslinked cells) and debris were pelleted at 12,000xg for 15 mins at 4°C. The supernatant containing chromatin was pre-cleared by adding 20µL washed Protein-A dynabeads (as above, resuspended in ChIP dilution buffer: 67mM Tris-HCl, 100mM NaCl, 5mM EDTA, 0.33% SDS, 1.67% Triton-X) and incubated at 4°C with rotation for at least 3 hours. 10µL of the pre-cleared chromatin was set aside as 5% input. For each reChIP, 200µL of pre-cleared chromatin ($1x10^6$ cells) was made up to 500µL in a dilution buffer to get a final concentration of 67mM Tris-HCl, 100mM NaCl, 5mM EDTA, 0.33% SDS, 1.67% Triton-X, 0.1µM PMSF, 1x protease inhibitors (reChIP buffer). 20µL Protein-A dynabeads were pre-incubated (as above) with either 1µg H3K4me3 (CST, #9751) or H2AK119ub (CST, #8240) antibodies or Rabbit IgG (Invitrogen,

#02-6102), then resuspended in the 500μL diluted chromatin. The first immunoprecipitation was performed overnight at 4°C with rotation, antibody-chromatin complexes were then washed three times in 500μL low-salt buffer (as above), three times in 500μL high-salt buffer (as above), twice in 200μL LiCl buffer (as above) and twice in 500μL 10mM TrisCl pH7.5 on ice. Complexes were eluted in 100μL ChIP elution buffer (as above) supplemented with 0.1μM PMSF and 1x protease inhibitors for 37°C for 30 mins on a thermomixer at 300RPM, of which 10μL were put aside as a 10% single ChIP control. To dilute the SDS, the eluate volume was increased to 300μL with reChIP buffer and purified using Amicon Ultra-0.5ml 3kDa filter columns (Millipore, #UFC5003) according to the manufacturer's instructions, with chromatin being additionally washed twice with 400μL ChIP dilution buffer. Purified chromatin was made up to 500μL in reChIP buffer for the second immunoprecipitation, which was performed using the alternate antibody or IgG control bead conjugates overnight at 4°C with rotation. Complexes were washed (as above) and resuspended in 100μL ChIP elution buffer. Samples were treated with 2μL RNase A and 2μL Proteinase K, then reversed crosslinked (as above). DNA was purified using the Monarch PCR & DNA Cleanup Kit (NEB) and eluted in 25μL $H_2O$ for qPCR and library preparation for sequencing.

**Cleavage under targets and release using nuclease (CUT&Run)**

NCI-H661 cells were harvested and counted after staining cells with 0.4% Trypan blue where 500,000 live cells were used per CUT&Run. Cells were washed with ice cold cell wash buffer (150mM NaCl, 20mM HEPES in $H_2O$) and kept on ice. 11uL Concanavalin A beads (Epicypher) were washed twice with 100uL cold bead activation buffer (20mM HEPES, 50uL KCl, 1mM $CaCl_2$, 1mM $MnCl_2$ in $H_2O$) and then resuspended in 11uL of the same buffer on ice. Cells were washed twice with NE buffer (20mM HEPES, 10mM KCl, 0.1% Triton X, 20% glycerol in $H_2O$) to release nuclei with gentle pipetting and 300xg spins at 4C in between for 5 mins each. Nuclei were resuspended in 100uL of cold cell wash buffer and then incubated with 10uL of washed beads to allow for nuclei-bead adsorption for 10 mins at room temperature. Beads were then resuspended in 50uL of cold antibody buffer (20mM HEPES, 150mM NaCl, 0.5mM Spermidine, 0.05% digitonin, 2mM EDTA in $H_2O$) followed by the addition of 1ug of IgG controls (Mouse IgG (Invitrogen, #31903), Rabbit IgG (Invitrogen, #012-6102)) or antibodies (DPPA2 (Sigma, #MAB4356), DPPA4 (abcepta, #AP1438a, abcepta, #AP21202c, abcam, #ab154642)) (or 0.5ug for the H3K4me3 (CST, #9751) positive control) and incubation overnight at 4C on a nutator. Bead complexes were then washed twice with 200uL cold antibody buffer (same as above, but without EDTA) and then resuspended in 50uL of the same buffer. 1.5uL of Protein-A/G-MNase (CST) was added to each bead suspension, pipette mixed and incubated at room temperature for 10 mins. Beads were washed two times with cold 250uL of antibody buffer and resuspended in 50uL of the same buffer. 1uL of 100mM $CaCl_2$ was added to each bead suspension to activate the bound MNase followed by incubation at 4C for 1hr on a thermalcycler to facilitate chromatin digestion. 38uL of stop buffer (340mM NaCl, 20mM EDTA, 4mM EGTA, 50ug/mL RNase A, 50ug/mL glycogen in $H_2O$) containing 250pg of *E.coli* normalisation spike-in DNA (CST) was added to each suspension to inhibit MNase, degrade RNA and release digested chromatin into the supernatant by incubation at 37C for 10 mins in a thermalcycler. The supernatant containing digested chromatin was transferred to a new tube and treated with 1uL 20mg/mL proteinase K for 1hr at 37C. DNA was purified using a 1.8X bead cleanup (SergiLabs) and eluted in 30uL of $H_20$ for DNA library preparation.

**Assay for transposase accessible chromatin (ATAC-seq) and library preparation**

NCI-H661 cells were harvested and counted after staining cells with 0.4% Trypan blue where 500,000 live cells were used for each ATAC assay. Cells were washed with ice cold PBS once and then lysed in 500uL NE buffer (10mM Tris-HCl, 10mM NaCl, 3mM $MgCl_2$, 0.1% IGEPAL CA-

630, 0.1% Tween-20) on ice by gentle pipetting to isolate nuclei. 50uL of nuclei (50,000 cells), were then pelleted at 1500xg for 10mins at 4C followed by supernatant aspiration. Nuclei were gently resuspended directly in tagmentation mix (1.75uL Tn5 transposase (Illumina), 17uL TD buffer (Illumina), 21.25uL $H_2O$) on ice and then allowed to tagment for 37C for 30 mins in a thermalcycler. Tagmented DNA was then purified using Ampure XP beads at a 1.8X ratio and eluted in 21uL of $H_2O$ for library preparation as previously described using 15 cycles of PCR to incorporate i7/i5 indices followed by a final 1.8X Ampure XP bead cleanup and elution in 22uL of $H_2O$.

**DNA library preparation and sequencing**

For ChIP, reChIP and CUT&Run libraries were prepared using the NEBNext UltraII DNA Library Preparation Kit (NEB) according to the manufacturer's instruction. During library preparation the NEBNext Multiplex Oligos for Illumina (Index Primers Set 1-4) were used to enable library multiplexing. Individual and pooled libraries were quantified and quality controlled using the dsDNA high sensitivity (HS) Qubit assay (Invitrogen) on the Qubit 3.0 fluorometer (Invitrogen) and HSD1000 High Sensitivity DNA kit (Agilent) on the TapeStation 4150 (Agilent). Libraries were pooled and sequenced on the Illumina NextSeq500 or NextSeq2000 platform in either single-end (RNA-seq, ChIP-seq) or paired-end (ATAC-seq, CUT&Run) configurations with 75-150bp read lengths targeting 15-30x$10^6$ 75-100bp single-end (ChIP, reChIP, RNA-seq), 10-15x$10^6$ 100bp paired-end (CUT&Run), 50x$10^6$ 100bp paired-end (ATAC-seq) or 100x$10^6$ 150bp paired-end (WGBS) reads.

**Nucleosome reconstitution**

Nucleosomes were reconstituted as described previously[5]. DNA used to generate nucleosomes was the 147 bp Widom 601 sequence[6] prepared by annealing complementary single stranded 147 bp Ultramers™ (Integrated DNA Technologies) with the sequence 5'-ATCGAGAATCCCGGTGCCGAGGCCGCTCAATTGGTCGTAGACAGCTCTAGCA CCGCTTAAACGCACGTACGCGCTGTCCCCCGCGTTTTAACCGCCAAGGGGATTACTCCCTAGTCT CCAGGCACGTGTCAGATATATACATCCGAT-3'. To prepare nucleosomes, 10 µM recombinant human histone octamers (histonesource.com) were buffer exchanged into buffer A (2 M NaCl, 10 mM Tris-HCl pH 7.5, 1 mM EDTA, 5 mM DTT) using a 7 kDa MWCO Zeba desalting column (Thermo Fisher), combined at a 1:1.2 molar ratio with Widom 601 dsDNA in buffer A and incubated for 30 min at 4 °C. The reconstitution reaction was dialyzed in buffer A with decreasing salt concentration stepwise from 2.0 M, 1.0 M, 0.8 M, 0.6 M to 0.2 M NaCl with incubation time of 2–3 hrs with 0.8 M NaCl buffer dialysis being an overnight step using a 7 kDa MWCO Slide-A-Lyzer mini dialysis unit (Life Technologies). The nucleosome reconstitution was confirmed by electromobility shift assay using 6% native acrylamide gel, visualized with ethidium bromide and Coomassie blue. The reconstituted nucleosomes were stored at 4 °C and used within 4 weeks of production.

**Electrophoretic mobility shift assay (EMSA)**

For nucleosome-binding assays, 50nM of recombinant nucleosomes were incubated in assay buffer (20 mM Tris-HCl pH 7.4, 75 mM NaCl, 6% glycerol 0.005% NP-40, 0.5 mM $MgCl_2$) with 0, 125, 250, 500 or 1000 nM recombinant DPPA2-Myc/Flag (Origene, TP305441), His-DPPA4 (Origene, TP760278) or DPPA2-DPPA4 complex (mixed in equi-molar concentration) in 10 µL reactions. The reactions were performed at room temperature for 30 min and resolved in a 0.5x TBE 6% 37.5:1 acrylamide: bis-acrylamide gel for 120 min at 100 volts at room temperature. A DNA ladder (GeneRuler 50bp; Thermo Fisher) was included in the gels. The gel was stained with SYBR Gold (Thermo Fisher) and imaged on ChemiDoc (BioRad). Each assay was repeated three times. The bands were quantified with ImageJ (ver2.14.0). Percentage of shifted nucleosomes is calculated by measuring the intensities of bottom bands and fraction of loss of bands against the nucleosomes only band.

**Mass photometry**

Mass photometry experiments were performed on Refeyn TwoMP mass photometer (Refeyn). Microscope coverslips (Refeyn) and silicon well gaskets (Refeyn) were cleaned with 100% isopropanol and water and dried with compressed air. The gaskets were placed on top of the cleaned coverslips on the sample stage of the mass photometer as per manufacturer's instructions. All measurements were performed at least three times independently, in separate wells, in buffer containing 20 mM HEPES pH 8, 50 mM NaCl, 1 mM TCEP. Firstly, 10 µL buffer was pipetted into a gasket-well followed by focal point acquisition using the autofocus functionality in the Refeyn AcquireMP 2.3.1 software. Secondly, 10 µL of 100 nM of recombinant DPPA2-Myc/Flag (Origene, TP305441), His-DPPA4 (Origene, TP760278) or in complex (mixed in equi-molar concentration and diluted to 100nM just prior to measurement to avoid complex disassembly) was pipetted in the buffer-containing gasket-well and mixed to observe individual landing events below saturation levels. Mass photometry movies of 6000 frames were recorded from a 10.8 × 10.8 µm instrument field of view. Data were processed using the default pipeline on the Refeyn DiscoverMP 2.3.0 software. Individual particle contrasts from each video were converted to mass using a contrast-to-mass (C2M) calibration which was performed in the acquisition buffer. Data were plotted as normalised histograms with a bin width of 4.4 and fitted to a Gaussian peak. For the calibration, 10 µL of a 1:30 pre-diluted Glutamine Synthetase (CLENZ544-2, Bio-Scientific) was added to an acquisition gasket-well; the 57, 231, 460 kDa masses were used for a standard calibration curve in the DiscoverMP software. The experiments were repeated three times independently.

**Quantitative PCR**

qPCR analysis of purified ChIP DNA and RT cDNA were performed in technical duplicate for each primer pair using 2x SYBR mastermix (Applied Biosystems, #4385612) according to manufacturer's instructions in a 6-12uL reaction on CFX96 or CFX384 (Bio-Rad) instruments.

***In vivo* animal studies**

NCI-H1299 cells containing doxycycline inducible overexpression pCLIIPi-Flag-DPPA2-PGK-mVenus and pCLIIPi-V5-DPPA4-PGK-tagBFP vectors stably integrated into the genome were induced with $H_2O$ or 2µg/mL doxycycline for 72 hrs in culture. Cells were harvested and counted following 0.4% Trypan blue staining, wherein $5x10^6$ cells were prepared in 50uL of RPMI-1640 with 50uL Matrigel (Corning) per mouse (10 mice per condition) without serum. 100uL of cells were subcutaneously injected into the right flank of 5-7 week-old male Balb/c nude mice (ARC). Mice were placed onto normal chow or chow containing 600mg/kg doxycycline two days after to maintain induction (10 mice per condition). Tumour volume was determined by electronic callipers and body weight by electronic scales at least weekly until tumours reached $1200mm^3$ volume or lost more than 20% of their body weight, upon which mice were humanely euthanised and organs, as well as the tumour harvested and weighed according to ethics application #2023-06 approved by the Peter MacCallum Cancer Centre (PMCC) Animal Ethics Committee. This work was conducted with the Translational Research Centre (TRC) at PMCC.

**TCGA, ERM and GTEx multiomic analyses**

Clinical information as well as matched transcriptome (RNA-seq), DNA methylation (methylation array), copy number and mutation (WES) data were obtained from the static LinkedOmics data repository for 11,518 cancer patients spanning 32 cancer types[7] from the cancer genome atlas (TCGA) study. RNA-seq untransformed gene level read counts were also obtained for NSCLC (LUAD and LUSC) patients from the genomic data commons (GDC) database. Processed TCGA pan-cancer chromatin accessibility (ATAC-seq) data[8] were also obtained from the GDC database. For mutations (WES), any mutation (SNV, deletion, insertion) on the DPPA2/4 gene body was considered. For copy number (WES) we classified ≥ 3 gene copies as a gain, ≤ 1 copies as a loss, and 2 copies as diploid. For ATAC-seq we utilised pre-compiled pan-cancer peaks and normalised counts at these peak regions for DPPA2/4 gene promoters (DPPA2p1 (TGCT_15780), DPPA4p1 (TGCT_15782), DPPA4p2 (TGCT_15783), DPPA4p3 (TGCT_15784)) or putative enhancers (DPPA4e1 (TGCT_15785), DPPA4e2 (TGCT_15786))[8]. Cox proportional hazard modelling (CoxPH) of patient outcomes (overall survival) were performed using the survival (v3.2-14) R package, where patients in each cohort was subdivided by the co-expression of DPPA2 and/or DPPA4 as determined by RNA-seq (expression cut-off defined as the median of all DPPA2 or DPPA4 expressors (RSEM>0)). Kaplan-Meier overall survival plots of patient groups were done using the survminer (v0.4.9) R package. Correlations between DPPA2/4 gene expression and copy number, promoter methylation and chromatin accessibility were assessed using Pearson correlation with the stats (v4.1.2) R package whereas correlations with binary mutation calls per patient were done using biserial correlation with the polyserial (v0.8-1) R package, dot plots visualisation these correlations were plotted using the corrplot (v0.92) R package. To discern differentially expressed gene in DPPA2/4 co-expressing NSCLC (LUAD, LUSC) tumours compared with DPPA2/4 non-expressing tumours we used untransformed read counts and performed differential gene expression analysis using the edgeR (v3.36.0) R package using the glmFit/LRT regression workflow whilst accounting for tumour subtypes in the model design (~Subtype+Group), where differentially expressed genes were considered as having an absolute log2FoldChange ≥ 1 and FDR < 0.05. All gene were then ranked by decreasing log2FC and then input to a gene set enrichment analysis (GSEA) using the fgsea (v1.20.0) R package against molecular signatures database (MSigDb) pathways with particular focus on the C2 and C8 pathway subsets. Processed gene expression (RNA-seq) data for n=19616 normal adult tissue samples across 30 tissue types were obtained from the Genotype-Tissue Expression (GTEx) database. Gene expression (RNA-seq) and DNA methylation (WGBS) data for human cell lines established across development were obtained from the Epigenetic Roadmap (ERM) (https://egg2.wustl.edu/roadmap/web_portal/). WGBS ERM data in bigwig format were imported into R using the rtracklayer package (v1.54.0). Since data were originally aligned to the hg19 human genome reference, genomic coordinates in these files were lifted-over using the liftOver R package (v1.18.0) with the hg19toHg38 UCSC liftover chain file (https://hgdownload.cse.ucsc.edu/goldenpath/hg19/liftOver/). Processed data were averaged over the DPPA2/4 promoter (TSS+/-1500bp) using the EnrichedHeatmap (v1.24.0) R package (normalizeToMatrix). All boxplots, barplots and scatter plots were plotted using the ggplot2 (v3.4.4) R package.

**NGS data pre-processing**

Single-end and paired-end reads in fastqs were trimmed and filtered for quality (phred33 score > 20) and length (>20bp) using TrimGalore (https://github.com/FelixKrueger/TrimGalore) v0.6.6 in single-end or paired-end mode (valid read pairs were only kept). Trimmed and quality filtered reads were then aligned to the hg38 human genome GRCh38.p13 (and for spike-in ChIP were also aligned to the mm10 mouse genome GRCm38.p6 separately) using bwa-mem[9] (bwa v0.7.13) with default parameters for ChIP-seq, CUT&Run, WGBS and ATAC-seq data. For ChIP-seq, CUT&Run and ATAC-seq, alignments were then converted to the bam format and indexed using samtools v1.9[10] . Duplicate alignments were marked with using MarkDuplicates (picard v2.6.0, https://broadinstitute.github.io/picard/) and re-indexed with samtools. For ATAC-seq data we additionally shifted reads by -4bp on the + strand and +5 bp on the -strand to account for Tn5 insertions using alignmentSieve (DeepTools v3.5.0)[11] . For spike-in ChIP data, we used the xenofilteR (v1.6)[12] R package to split human and mouse aligned reads. bigWig files containing CPM/bp normalised coverage values for each sample were derived from duplicate marked bam files using bamCoverage (DeepTools v3.5.0) where single-end data were extended to 147bp and paired-end data were not extended (relying instead on the true fragment length determined by read-pairs).

For WGBS data following read trimming as above, Bismark (v0.22.3)[13] was used to align reads to a bisulfite converted hg38 (GRCh38.p13) genome using bowtie2 internally, followed by read deduplication and methylated/unmethylated read counting. The resultant methylation bedgraph was converted to the bigwig format using bedGraphToBigWig (UCSC.utils (v1.3.1) R package).

For RNA-seq data following read trimming as above, reads were aligned using the splice-aware aligner STAR (v2.7.5b)[14] and gene counts were made using featureCounts (subread v2.0.6)[15] with human GENCODE v36 gene annotations.

**Peak calling, annotation, differential testing (ChIP-seq, reChIP-seq, CUT&Run, ATAC-seq)**

Peak calling for histone modification ChIPs (H3K4me3, H3K4me1, H3K27ac, H3K27me3, H2AK119ub, H3K9me3, H3K36me3 and H3K36me3) as well as reChIPs were performed separately for independent replicates using EPIC2 (v0.0.52)[16] (--bin-size 100 –gaps-allowed 1 –fragment-size 147 -- false-discovery-rate-cutoff 0.05) using matched genomic inputs as the control. For DPPA2, DPPA4, SUZ12 ChIP we used MACS2[17] (v2.2.7.1) to call narrow peaks, using matched genomic inputs as the control (-B --nomodel --extsize 147 --SPMR -q 0.05). For CUT&Run and ATAC-seq we also used MACS2 to call peaks, except with options more suitable for paired-end data (-f BAMPE -q 0.05) using matched IgG's as the control for CUT&Run and no controls for ATAC-seq. Downstream data analyses were conducted using R (v4.1.2).

Peaks were filtered (log2FC >1 and FDR < 0.05) and regions overlapping blacklisted regions from the hg38 ENCODE blacklist were excluded[18] . To derive commonly enriched regions between replicates we retained and merged peaks with >25% overlap in replicates using the GenomicRanges R package (findOverlaps, Reduce and union) (v1.46.1). For reChIP analyses H3K4me3-H2AK119ub and H2AK119ub-H3K4me3 peaks were also merged as above to derive reciprocally bivalent regions. All consensus peak sets following merging were annotated using HOMER (annotatePeaks.pl) (v4.11)[19] which by default denotes promoter-TSS regions as 1kb upstream and 100bp downstream of transcription start sites (TSS). Promoter-TSS regions were re-defined as the region spanning 1.5kb upstream and 1.5kb downstream of TSS's.

For differential enrichment analyses a consensus peak set was first derived using the GenomicRanges R package (Reduce and union) where atleast >50% overlap between peaks of replicates had to be observed, form which a final union was taken. Reads were counted at consensus peaks using the csaw R package[20] (regionCounts) (v1.28.0). Normalisation factors for each sample were pre-computed by binning the genome into 10kb windows, generating counts for each 10kb window and performing trimmed mean of M-values (TMM) normalisation using csaw (normFactors) to adjust for compositional biases. Differential enrichment was then performed using edgeR (estimateDisp>glmFit>glmLRT) (v3.36.0) in which doxycycline conditions were contrasted with their matched vehicle control per cell line. Since experiments were replicated in complete batches we accounted for this batch-effect in design models for differential testing (~Batch+Treatment). Benjamini-Hochberg p-value corrections for multiple testing were applied to these contrasts. Significant differential enrichment was defined as a region that had an absolute log2FC ≥ 0.5 and FDR < 0.05.

CpG island and ENCODE cis-regulatory element overlaps were performed using the GenomicRanges R package (findOverlaps) and were obtained for the hg38 genome using the UCSC table browser. Odds ratio testing to test for representation of sets of regions were performed using the fmsb (v0.7.3) R package, of which resultant p-values were corrected for multiple comparisons using the Benjamini-Hochberg method.

**Differential gene expression analysis (RNA-seq)**

Differential testing for RNA-seq data were conducted on unstransformed gene counts (filtered for all expressed genes (>1 count in any sample) using the edgeR R package (estimateDisp>glmFit>glmLRT). For siRNA knockdown and shRNA knockdown comparisons data were generated in complete batches we accounted for this batch-effect in design models for differential testing (~Batch+Treatment) relative to a treatment control. For shRNA knockdowns we noted a prominent doxycycline-induced transcriptional program in the above comparison that was later accounted for using an additional design model (~shRNA+Treatment+shRNA:Treatment) using the shREN (control shRNA) condition to model the global effect of doxycycline on transcription. For principal component analyses we utilised batch-corrected counts from edgeR (removeBatchEffect).For all comparisons differentially expressed genes were considered as having an absolute log2FoldChange ≥ 1 and FDR < 0.05. All boxplots, barplots and scatter plots were plotted using the ggplot2[21] (v3.4.4) R package.

**Differential methylation analysis (WGBS)**

Bismark coverage files reporting per-CpG methylation counts were imported to R for use in differential methylation analyses with edgeR (readBismark2DGE). We retained those CpGs that had a read count coverage of atleast 10 in all samples and that were not solely hypomethylated or hypermethylated for differential testing. We then performed differential testing between treatments and controls, accounted for batch-effects in the design model (~Batch+Treatment) using a modified design matrix that accounts for both methylated and unmethylated counts (modelMatrixMeth) using edgeR (estimateDisp>glmFit>glmLRT). Differentially methylated CpGs were those with an FDR < 0.05. Scatter plots were plotted using the ggplot2 (v3.4.4) R package

**Genomic enrichment heatmaps and trackplots (ChIP-seq, reChIP-seq, CUT&Run, WGBS)**

To generate genomic enrichment heatmaps and trackplots bigWigs containing CPM/bp normalised read densities were imported to R using the rtracklayer package (import.bw) (v1.54.0)[22] . For heatmaps, each peak region was first extended to 5kb upstream and downstream and then split into 100 equally sized bins using the GenomicRanges R package (resize) (v1.46.1). The average CPM/bp was calculated for each bin for each ChIP using the EnrichedHeatmap R package (normalizeToMatrix) (v1.24.0). Bins with values surpassing the 99th percentile of all bins within each ChIP were masked (i.e. assigned the 99th percentile value) to eliminate extreme outliers from affecting colour scales. Each bin was then scaled relative to the highest value (so values range between 0-1 and represent the relative enrichment of signal across all regions), except for ChIPs where treatments were performed (e.g. vehicle control and doxycycline) which were left unscaled and WGBS data which were already between 0-1 (methylation fraction) and did not need scaling. Enriched heatmaps were then plotted using the same package (EnrichedHeatmap), with the average bin value plotted as continuous curves atop each heatmap. Genomic track plots were plotted using the rtracklayer (v1.54.0) and Gviz R

[23] packages (v1.38.4) with no prior scaling and represent either CPM/bp or methylation fractions. CpG island annotations and ENCODE cis-regulatory elements for the hg38 genome were retrieved from the UCSC table browser.

**Gene ontology**

The enrichment of gene ontologies across subclasses of differentially expressed or enriched genes were determined using the clusterProfiler R package (v4.2.2). Gene symbols were first converted to entrez id's using the biomaRt[24] R package (v2.50.3) and were input alongside a background list of all expressed genes to clusterProfiler (compareCluster) against the Gene Ontology (GO) Biological Processes (BP) database. Significantly enriched GO terms were those with a Benjamini-Hochberg (BH) corrected p-value < 0.05, had atleast 10 genes present in the pathway and a gene ratio (genes in subclass/genes in pathway) > 0.01. Representative pathways were plotted using the ggplot2 R package (v3.3.5).

**CG-content and motif analysis**

For all sequence based analyses, randomised sequences were modelled off the width and re-sampled based on the GC-content of peak regions of interest (made with the regioneR R package (createRandomRegions)[25] (v1.26.1) and nullranges R package (matchRanges R package[26], providing GC-content as a covariate) (v1.0.1)) and were used as the background for downstream analyses. CG content was determined for different region sets as well as background sequences above using the Biostrings R package (oligonucleotideFrequency) (v2.62.0) by first calculating all oligonucleotide frequencies and then by summing C and G frequencies. All dinucleotide frequencies were calculated using monaLisa[27] (plotBinDiagnostics) and then GC/CG dinucleotide frequencies were summed. These data were plotted using ggplot2, and the significance of comparisons were determined using pairwise t-tests followed by BH-adjustments of p-values to account for multiple comparisons.

De novo motif discovery was undertaken for DPPA2+4 consensus binding regions following resizing of these regions as well as matched background regions to 1kb (median length of regions) using HOMER (v4.11, findMotifsGenome.pl).

Enrichments for known transcription factor binding motifs in differential peak subclasses were calculated using the monaLisa R package (v1.0.0) following resizing of all regions to 4kb (median length of all differential/background regions). Position weight matrices for transcription factor binding sites in vertebrates were retrieved from the JASPAR2020 database. Binned motif enrichment for region sets were then conducted in monaLisa (calcBinnedMotifEnrR) where significant enrichments were those with a BH-adjusted p-value < 0.05 and log2 Fold Change over background sequences > 0.25. Motif heatmaps were also plotted using monaLisa (plotMotifHeatmaps).

**Chromatin state discovery (ChromHMM)**

H3K4me3, H3K4me1, H3K27ac, H3K27me3, H2AK119ub, H3K9me3, H3K36me2 and H3K36me3 ChIP bam files were first converted to the bed format using bedtools (bamtobed) (v2.27.1). bed files were then partitioned into 200bp bins and then binarized for the determination of bin-specific enrichments (input as the control) using ChromHMM (BinarizeBed) (v1.24)[28]. To incorporate WGBS DNA methylation data into the model we computed methylation fractions in same bins using Bismark coverage files imported via the bsseq (v1.30.0) R package, and binarized bins such that enrichment was indicated as >0.5

methylation fraction. Hidden Markov Models were then used to discover chromatin states across these genomic bins using ChromHMM (LearnModel) using a 17-state model. Segment bed files containing chromatin state annotations were then overlapped with DPPA2+4 consensus peaks, where each peak was then re-assigned to the chromatin state with the highest degree of overlap using the GenomicRanges R package (findOverlaps and pintersect) (v1.46.1). Heatmaps containing emission probabilities, transition probabilities, TSS enrichments and annotation overlaps from the ChromHMM model were then plotted using the ComplexHeatmap R package (v2.10.0).

For the integrative ChromHMM model with 27 NSCLC cell lines we performed a joint 14-state ChromHMM model for those marks which were available via DBTSS[29] (H3K4me3, H3K4me1, H3K27ac, H3K27me3, H3K9me3, and H3K36me3) without DNA methylation. To summarise chromatin states at sets of regions, we called the most frequent ChromHMM state annotation per region. Transitioning states were those defined as being in atleast 75% of all other NSCLC cell lines (n=26) for a given DPPA2+4 binding site and were different from the NCI-H661 chromatin state at that site.

**Software**

Plots were generated using R (v4.1.2/RStudio v2022.02.0+443) and edited in Inkscape. Schematic figures were made with BioRender.com with publishing licence agreement numbers RM25UDOLG3, UQ25UDOE0U and MC25UDOPHO.

**Supplemental Table 1: shRNA oligo sequences**

| Name | Forward | Reverse |
|---|---|---|
| REN.713 | tcgagaaggtatatTGCTGTTGACAGTGAGCGCAGGAATTATAATGCTTATCTATAGTGAAGCCACAGATGTATAGATAAGCATTATAATTCCTATGCCTACTGCCTCGG | aattCCGAGGCAGTAGGCATAGGAATTATAATGCTTATCTATACATCTGTGGCTTCACTATAGATAAGCATTATAATTCCTGCGCTCACTGTCAACAGCAatataccttc |
| DPPA2.760 | tcgagaaggtatatTGCTGTTGACAGTGAGCGAACCAATACAGTTGAAGTGATATAGTGAAGCCACAGATGTATATCACTTCAACTGTATTGGTCTGCCTACTGCCTCGG | aattCCGAGGCAGTAGGCAGACCAATACAGTTGAAGTGATATACATCTGTGGCTTCACTATATCACTTCAACTGTATTGGTTCGCTCACTGTCAACAGCAatataccttc |
| DPPA4.1993 | tcgagaaggtatatTGCTGTTGACAGTGAGCGCTAAGATGTGTATGTAAAATAATAGTGAAGCCACAGATGTATTATTTTACATACACATCTTATTGCCTACTGCCTCGG | aattCCGAGGCAGTAGGCAATAAGATGTGTATGTAAAATAATACATCTGTGGCTTCACTATTATTTTACATACACATCTTAGCGCTCACTGTCAACAGCAatataccttc |

**Supplemental Table 2 siRNA sequences**

| SMARTpool | Individual siRNA in pool | Target sequence (5’-3’) |
|---|---|---|

| | | |
|---|---|---|
| ON-TARGETplus Non-targeting Control Pool<br><br>D-001810-10-05 | ON-TARGETplus Non-targeting #1<br><br>D-001810-01 | UGGUUUACAUGUCGACUAA |
| | ON-TARGETplus Non-targeting #2<br><br>D-001810-02 | UGGUUUACAUGUUGUGUGA |
| | ON-TARGETplus Non-targeting #3<br><br>D-001810-03 | UGGUUUACAUGUUUUCUGA |
| | ON-TARGETplus Non-targeting #4<br><br>D-001810-04 | UGGUUUACAUGUUUUCCUA |
| ON-TARGETplus Human DPPA2 (151871) siRNA SMARTpool<br><br>L-018977-01-0005 | ON-TARGETplus SMARTpool DPPA2<br><br>J-018977-09 | CAGUUAAAGAUGACGCAAA |
| | ON-TARGETplus SMARTpool DPPA2<br><br>J-018977-10 | CAAAUGGAACCAAGCGUUU |
| | ON-TARGETplus SMARTpool DPPA2<br><br>J-018977-11 | CGGGACACUUUGCGGGACU |
| | ON-TARGETplus SMARTpool DPPA2<br><br>J-018977-12 | CGACUGUGCUAAGAGGAAU |
| ON-TARGETplus Human DPPA4 (55211) siRNA SMARTpool<br><br>L-020766-01-0005 | ON-TARGETplus SMARTpool DPPA4<br><br>J-020766-09 | CCACAGAGAAGUCGAGGGA |
| | ON-TARGETplus SMARTpool DPPA4<br><br>J-020766-10 | GGUGUGUGGUCCAUGGGAA |

| | | |
|---|---|---|
| | ON-TARGETplus SMARTpool DPPA4<br><br>J-020766-11 | CCGAUUCUCCAUAAAUAAA |
| | ON-TARGETplus SMARTpool DPPA4<br><br>J-020766-12 | GUAAAGUGCUCUGCCCUAA |

**Appendix 1: ORFs:**

StrepII-Flag-DPPA2 fragment (NM_138815):

TCGAAAGGATCCTTAATACGACTCACTATAGGGAGACCCAAGCTGGCTAGCCACCATGGATTATAA
AGATGATGATGATAAAGGGTCGGCCGCCAGCTGGAGCCACCCTCAGTTCGAGAAGGGAGGAGGA
AGCGGCGGAGGCAGCGGAGGAGGAAGCTGGAGCCACCCGCAGTTCGAGAAAGGAGCTAGATC
AGAGAACCTGTACTTCCAATCCATGTCAGATGCAAATTTGGATAGCAGCAAGAAGAATTTCTTGGAG
GGGGAAGTAGATGATGAGGAAAGTGTGATTTTGACACTGGTGCCAGTTAAAGATGACGCAAATATG
GAACAAATGGAACCAAGCGTTTCTTCAACTTCTGATGTCAAACTGGAGAAGCCTAAGAAATACAATC
CAGGTCATCTACTTCAAACAAATGAGCAATTTACAGCTCCACAAAAAGCTAGATGCAAAATACCAGC
CCTTCCCTTGCCGACCATTTTGCCTCCCATTAATAAGGTGTGTCGGGACACTTTGCGGGACTGGTG
TCAACAACTCGGTTTGAGTACTAATGGCAAGAAAATCGAAGTTTATCTGAGGCTTCATAGGCATGCTT
ACCCTGAACAACGGCAAGATATGCCTGAAATGTCACAAGAGACCAGATTACAGCGATGTTCGAGG
AAACGCAAGGCAGTGACCAAGAGAGCAAGGCTTCAGAGAAGTTATGAGATGAATGAGAGAGCAGA
AGAGACCAATACAGTTGAAGTGATAACTTCAGCACCGGGAGCCATGTTGGCATCATGGGCAAGAAT
TGCTGCAAGAGCTGTTCAGCCTAAGGCTTTGAATTCATGTTCCATTCCTGTTTCTGTTGAGGCCTTTTT
GATGCAAGCCTCTGGCGTCAGGTGGTGTGTGGTCCATGGCAGACTTCTCTCGGCAGACACAAAGG
GTTGGGTACGCCTGCAGTTTCATGCAGGTCAGGCCTGGGTGCCTACCACTCACAGGAGGATGATT
TCTCTCTTCTTGTTACCTGCCTGCATTTTCCCATCCCCAGGCATAGAAGATAATATGTTATGCCCCGA
CTGTGCTAAGAGGAATAAGAAGATGATGAAAAGATTAATGACAGTAGAGAAGTAAGGCGCGGCCGG
CCAGCCG

StrepII-V5-DPPA4 fragment (NM_018189):

TCGAAAGGATCCTTAATACGACTCACTATAGGGAGACCCAAGCTGGCTAGCCACCATGGGTAAGC
CTATCCCTAACCCTCTCCTCGGTCTCGATTCTACGGGGTCGGCCGCCAGCTGGAGCCACCCTCA
GTTCGAGAAGGGAGGAGGAAGCGGCGGAGGCAGCGGAGGAGGAAGCTGGAGCCACCCGCAGT
TCGAGAAAGGAGCTAGATCAGAGAACCTGTACTTCCAATCCATGTTGCGAGGCTCCGCTTCTTCTA
CAAGTATGGAGAAGGCAAAAGGCAAGGAGTGGACCTCCACAGAGAAGTCGAGGGAAGAGGATCA
GCAGGCTTCTAATCAACCAAATTCAATTGCTTTGCCAGGAACATCAGCAAAGAGAACCAAAGAAAAA
ATGTCTGTCAAAGGCAGTAAAGTGCTCTGCCCTAAGAAAAAGGCAGAGCACACTGACAACCCCAG
ACCTCAGAAGAAGATACCAATCCCTCCATTACCTTCTAAACTGCCACCTGTTAATCTGATTCACCGG
GACATTCTGCGGGCCTGGTGCCAACAATTGAAGCTGAGCTCCAAAGGCCAGAAATTGGATGCATA
TAAGCGCCTGTGTGCCTTTGCCTACCCAAATCAAAAGGATTTTCCTAGCACAGCAAAAGAGGCCAA
AATCCGGAAATCATTGCAAAAAAAATTAAAGGTGGAAAAGGGGGAAACGTCCCTGCAAAGTTCTGA
GACACATCCTCCTGAAGTGGCTCTTCCTCCTGTGGGGGAGCCGCCTGCCCTGGAAAATTCCACTG
CTCTCCTTGAGGGAGTTAATACAGTTGTGGTGACAACTTCTGCCCCAGAGGCTTTGCTGGCCTCCT
GGGCGAGAATTTCAGCCAGGGCGAGGACACCAGAGGCAGTGGAATCTCCACAAGAGGCCTCTG

GTGTCAGGTGGTGTGTGGTCCATGGGAAAAGTCTCCCTGCAGACACAGATGGTTGGGTTCACCTG
CAGTTTCATGCTGGTCAAGCCTGGGTTCCAGAAAAGCAAGAAGGGAGAGTGAGTGCACTCTTCTTG
CTTCCTGCCTCCAATTTTCCACCCCCGCACCTTGAAGACAATATGTTGTGCCCCAAATGTGTTCACA
GGAACAAGGTCTTAATAAAAAGCCTCCAATGGGAATAAGGCGCGGCCGGCCAGCCG

**Supplemental Information References**


1. Eckersley-Maslin, M. A. *et al.* Epigenetic priming by Dppa2 and 4 in pluripotency facilitates multi-lineage commitment. *Nat Struct Mol Biol* **27**, 696–705 (2020).

2. Fellmann, C. *et al.* An Optimized microRNA Backbone for Effective Single-Copy RNAi. *Cell Rep.* **5**, 1704–1713 (2013).

3. Narita, M. *et al.* A Novel Role for High-Mobility Group A Proteins in Cellular Senescence and Heterochromatin Formation. *Cell* **126**, 503–514 (2006).

4. Moudgil, A. *et al.* Self-Reporting Transposons Enable Simultaneous Readout of Gene Expression and Transcription Factor Binding in Single Cells. *Cell* **182**, 992-1008.e21 (2020).

5. Tan, W. *et al.* Preparation and purification of mono-ubiquitinated proteins using Avi-tagged ubiquitin. *PLoS ONE* **15**, e0229000 (2020).

6. Lowary, P. T. & Widom, J. New DNA sequence rules for high affinity binding to histone octamer and sequence-directed nucleosome positioning. *J. Mol. Biol.* **276**, 19–42 (1998).

7. Vasaikar, S. V., Straub, P., Wang, J. & Zhang, B. LinkedOmics: analyzing multi-omics data within and across 32 cancer types. *Nucleic Acids Res.* **46**, gkx1090- (2017).

8. Corces, M. R. *et al.* The chromatin accessibility landscape of primary human cancers. *Science* **362**, (2018).

9. Li, H. & Durbin, R. Fast and accurate short read alignment with Burrows–Wheeler transform. *Bioinformatics* **25**, 1754–1760 (2009).

10. Li, H. *et al.* The Sequence Alignment/Map format and SAMtools. *Bioinformatics* **25**, 2078–2079 (2009).

11. Ramírez, F., Dündar, F., Diehl, S., Grüning, B. A. & Manke, T. deepTools: a flexible platform for exploring deep-sequencing data. *Nucleic acids Res.* **42**, W187-91 (2014).

12. Kluin, R. J. C. *et al.* XenofilteR: computational deconvolution of mouse and human reads in tumor xenograft sequence data. *Bmc Bioinformatics* **19**, 366 (2018).

13. Krueger, F. & Andrews, S. R. Bismark: a flexible aligner and methylation caller for Bisulfite-Seq applications. *Bioinformatics* **27**, 1571–2 (2011).

14. Dobin, A. *et al.* STAR: ultrafast universal RNA-seq aligner. *Bioinformatics* **29**, 15–21 (2012).

15. Liao, Y., Smyth, G. K. & Shi, W. The Subread aligner: fast, accurate and scalable read mapping by seed-and-vote. *Nucleic Acids Res.* **41**, e108–e108 (2013).

16. Stovner, E. B. & Sætrom, P. epic2 efficiently finds diffuse domains in ChIP-seq data. *Bioinformatics* **35**, btz232 (2019).

17. Zhang, Y. *et al.* Model-based analysis of ChIP-Seq (MACS). *Genome Biol.* **9**, R137 (2008).

18. Amemiya, H. M., Kundaje, A. & Boyle, A. P. The ENCODE Blacklist: Identification of Problematic Regions of the Genome. *Sci. Rep.* **9**, 9354 (2019).

19. Heinz, S. *et al.* Simple Combinations of Lineage-Determining Transcription Factors Prime cis-Regulatory Elements Required for Macrophage and B Cell Identities. *Mol. Cell* **38**, 576–589 (2010).

20. Lun, A. T. L. & Smyth, G. K. csaw: a Bioconductor package for differential binding analysis of ChIP-seq data using sliding windows. *Nucleic Acids Res.* **44**, e45–e45 (2016).

21. Valero-Mora, P. M. ggplot2: Elegant Graphics for Data Analysis. *J. Stat. Softw.* **35**, (2010).

22. Lawrence, M., Gentleman, R. & Carey, V. rtracklayer: an R package for interfacing with genome browsers. *Bioinformatics* **25**, 1841–1842 (2009).

23. Yu, G., Wang, L.-G., Han, Y. & He, Q.-Y. clusterProfiler: an R Package for Comparing Biological Themes Among Gene Clusters. *OMICS: A J. Integr. Biol.* **16**, 284–287 (2012).

24. Smedley, D. *et al.* BioMart – biological queries made easy. *BMC Genom.* **10**, 22–22 (2009).

25. Gel, B. *et al.* regioneR: an R/Bioconductor package for the association analysis of genomic regions based on permutation tests. *Bioinformatics* **32**, 289–291 (2016).

26. Davis, E. S. *et al.* matchRanges: generating null hypothesis genomic ranges via covariate-matched sampling. *Bioinformatics* **39**, btad197 (2023).

27. Machlab, D. *et al.* monaLisa: an R/Bioconductor package for identifying regulatory motifs. *Bioinformatics* **38**, 2624–2625 (2022).

28. Ernst, J. & Kellis, M. ChromHMM: automating chromatin-state discovery and characterization. *Nat. Methods* **9**, 215–216 (2012).

29. Suzuki, A. *et al.* DBTSS as an integrative platform for transcriptome, epigenome and genome sequence variation data. *Nucleic Acids Res.* **43**, D87–D91 (2015).
