## Supplemental Figures for "Embryonic stem cell factors DPPA2/4 facilitate a unique chromatin state in non-small cell lung cancer"

### Extended Figure 1, related to Figure 1

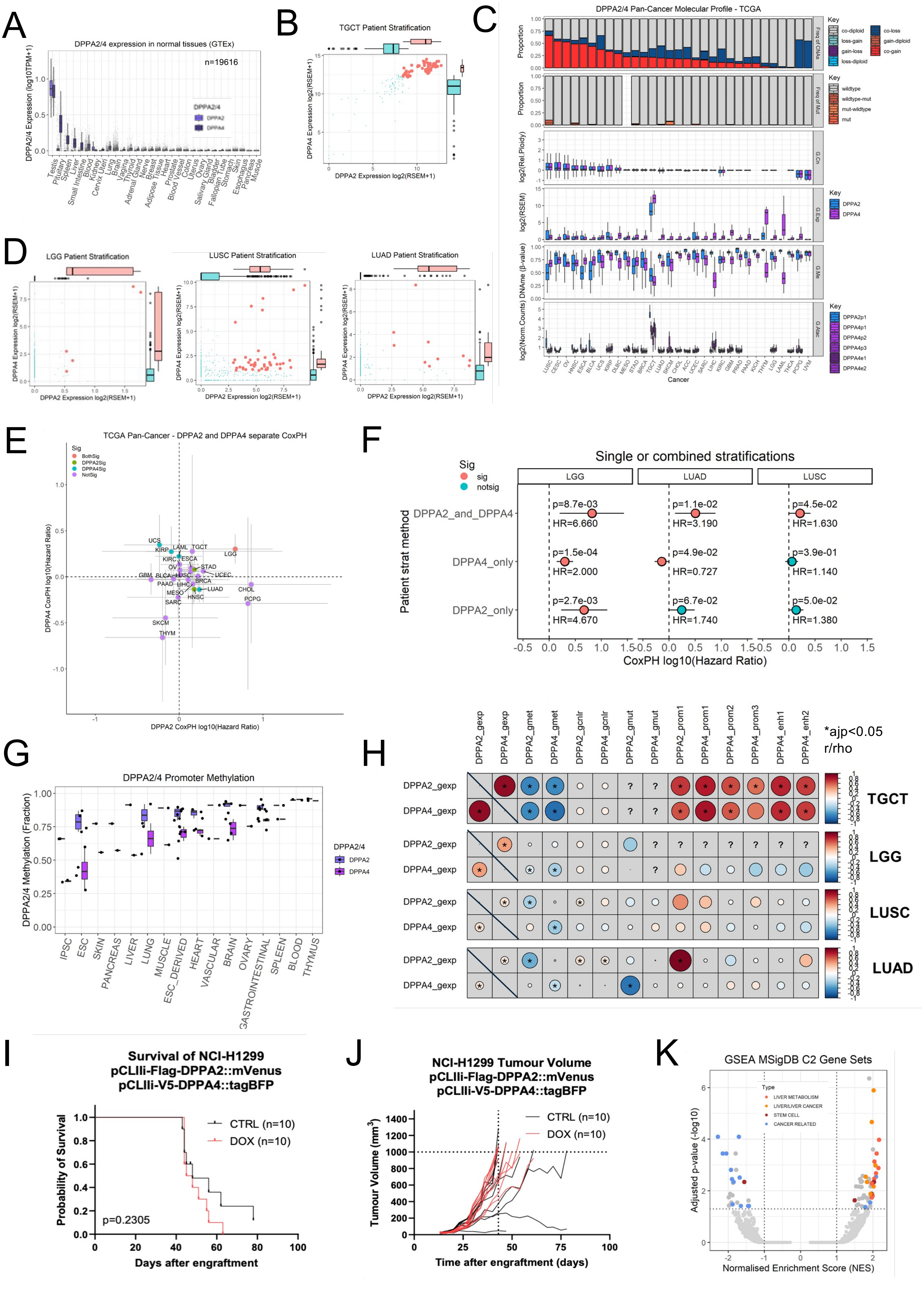

### Extended Figure 2, relating to Figure 2

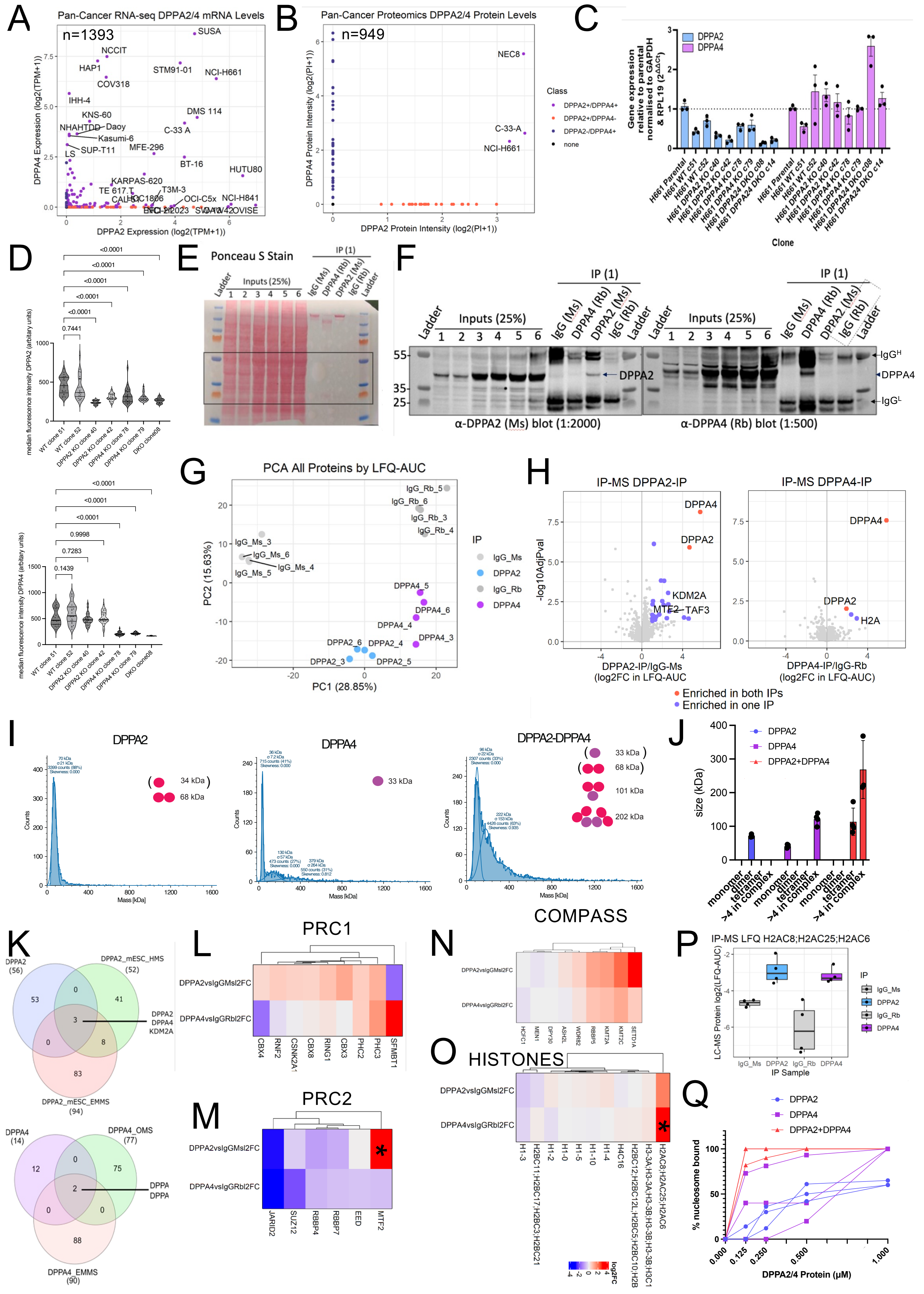

### Extended Figure 3, related to Figure 3

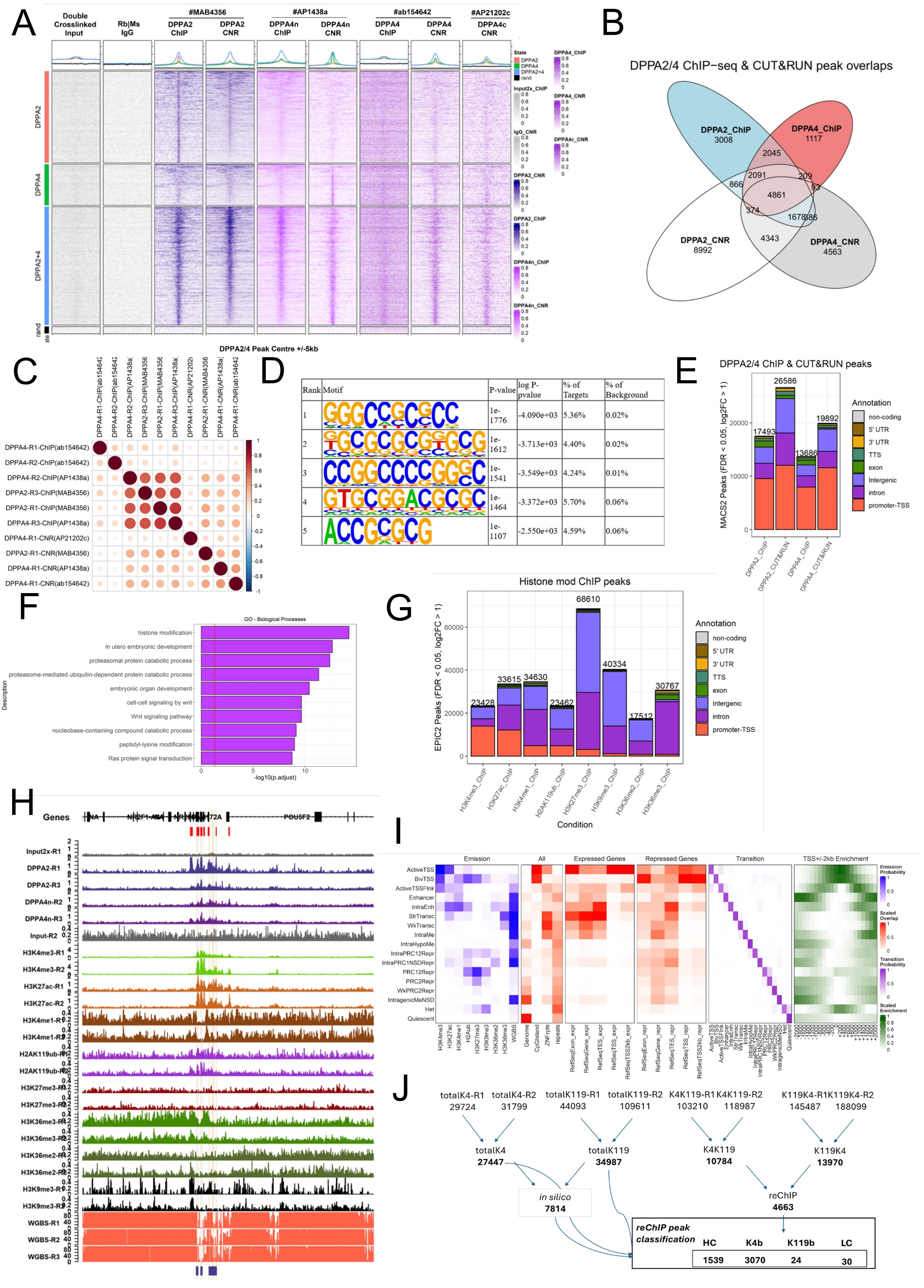

### Extended Figure 4, relating to Figure 4

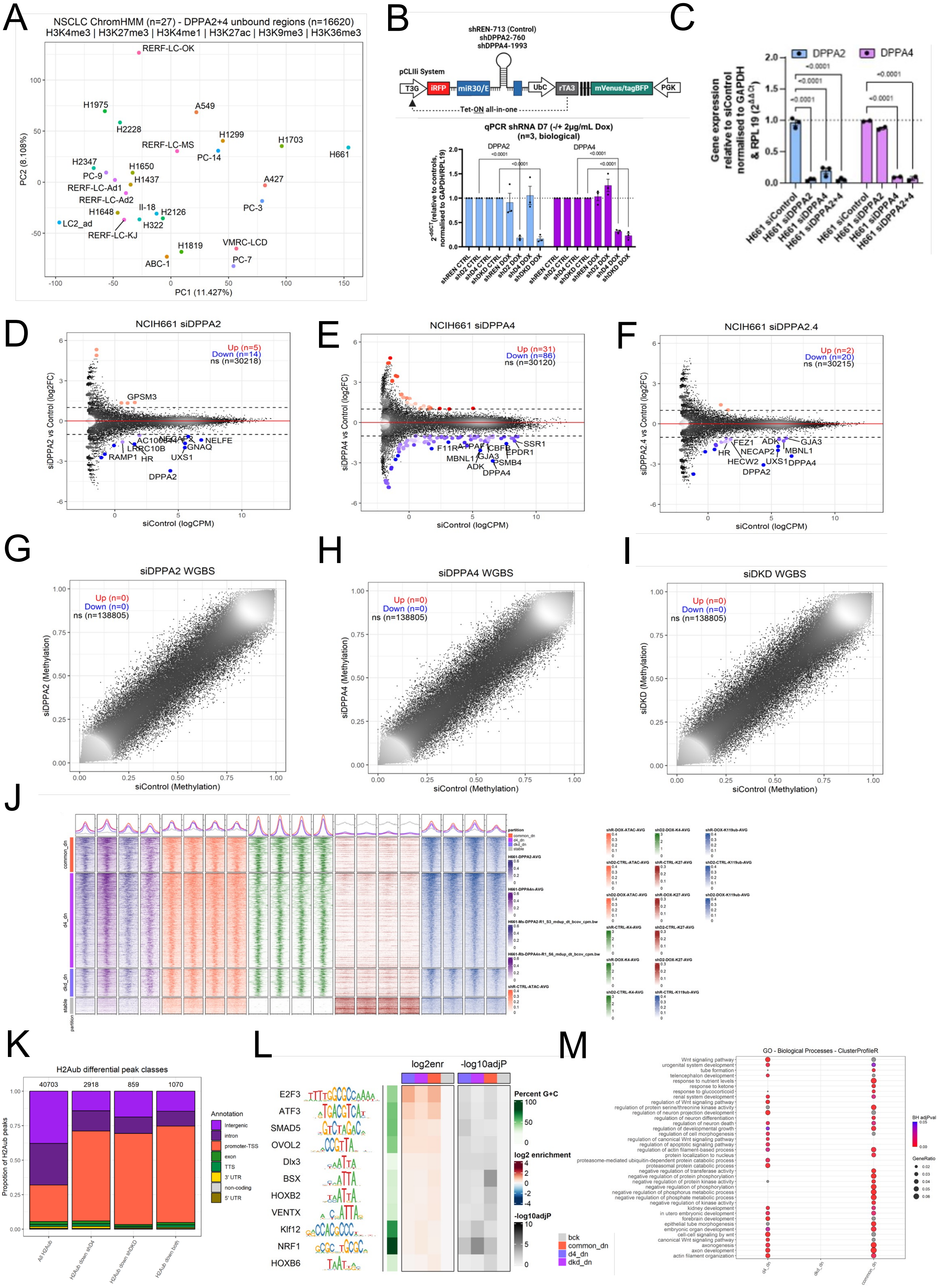

### Extended Figure 5, relating to Figure 5

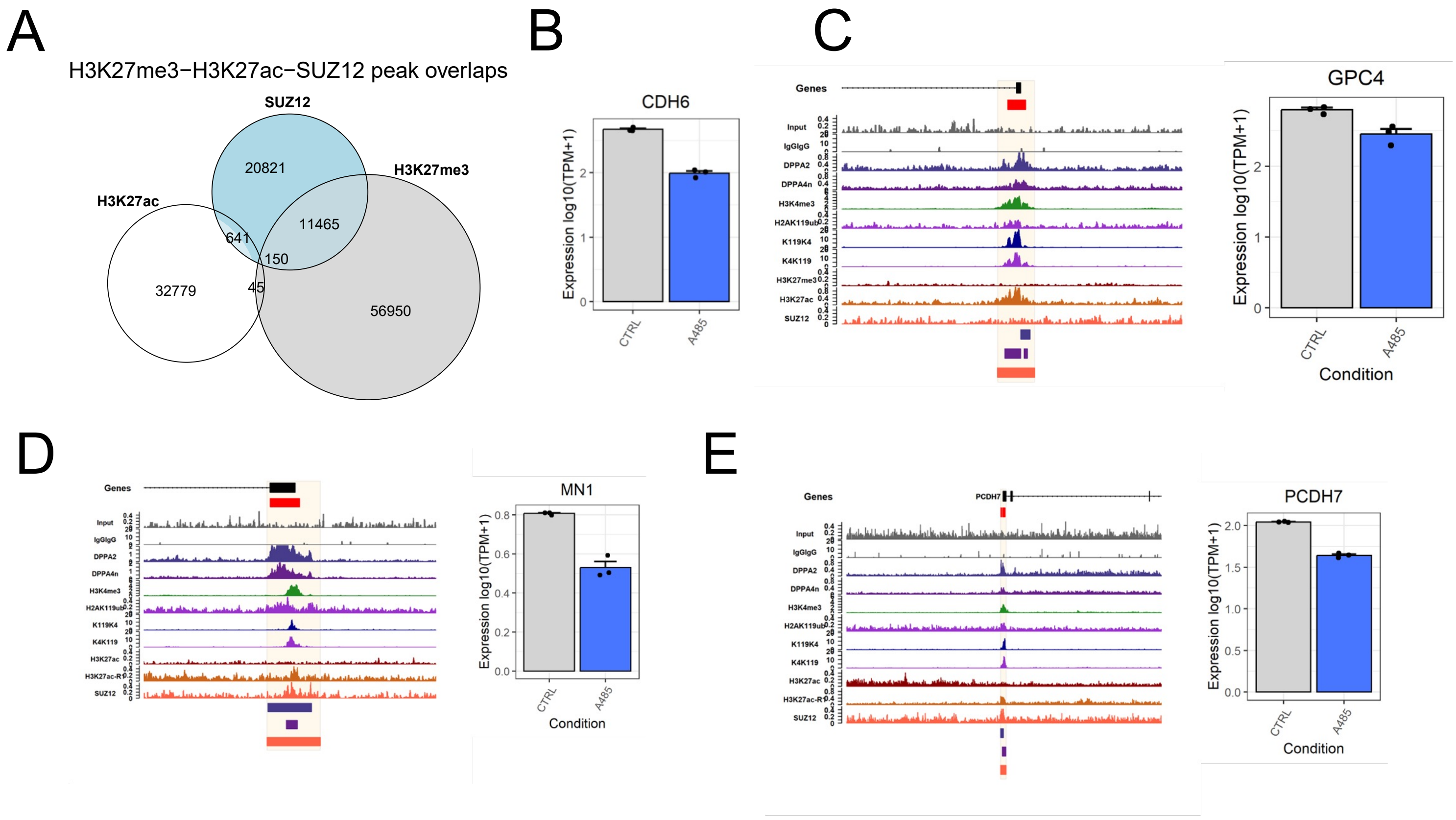
